# Angiotensin II–Driven Coronary Vasculopathy and Pressure-Overload Myocardial Remodeling Represent Distinct Vascular Phenotypes

**DOI:** 10.64898/2026.06.21.733633

**Authors:** Dzmitry Matsiukevich, David M. Ornitz

## Abstract

**Objective:** Chronic activation of the renin-angiotensin-aldosterone system (RAAS) promotes pathological remodeling of both myocardium and coronary arteries, yet the mechanisms that distinguish myocardial from vascular remodeling remain poorly defined. This study dissects the relative contributions of hemodynamic versus neurohumoral stress to cardiac remodeling, with emphasis on coronary vasculopathy and vascular smooth muscle cell (VSMC) plasticity.

**Methods:** Three murine models were used: transverse aortic constriction (TAC), angiotensin II (AngII) plus phenylephrine (AngII/PE), and high-dose angiotensin II (HD-AngII). Hemodynamics were assessed by catheterization at early and late time points. Histological and immunostaining analyses quantified interstitial and perivascular remodeling, including cardiomyocyte hypertrophy, interstitial and perivascular fibrosis, VSMC phenotype transitions, proliferation and quiescence markers, and neointimal and elastic lamina remodeling.

**Results:** After 28 days, all models exhibited diastolic dysfunction and myocardial fibrosis. Systolic pressure averaged ∼130 mmHg in both AngII models versus ∼200 mmHg in TAC. Despite lower pressure, myocardial fibrosis was greater in AngII/PE and HD-AngII models. While TAC induced uniform cardiomyocyte hypertrophy, hypertrophy in AngII models localized near fibrotic and perivascular regions. Increasing AngII dosage shifted remodeling from predominantly myocardial to predominantly vascular phenotypes, accompanied by VSMC dedifferentiation, proliferation, centripetal migration across the internal elastic lamina, neointima formation, elastic lamina disruption, and increased circulating desmosine, consistent with elastin degradation. AKT signaling was selectively increased in coronary VSMCs during this vasculopathic remodeling. Lineage-tracing analyses showed that Ang II–driven coronary neointima formation occurs beneath an intact endothelial monolayer and is composed predominantly of VSMC-derived cells, highlighting a VSMC-centric vasculopathy distinct from classic endothelium-initiated vascular remodeling.

**Conclusion:** Hemodynamic pressure overload and AngII-dominant neurohumoral stress drive distinct cardiac remodeling phenotypes: TAC primarily elicits uniform myocardial hypertrophy and interstitial fibrosis, whereas chronic AngII exposure preferentially promotes a VSMC-centric coronary vasculopathy with perivascular fibrosis and elastic lamina injury at lower pressure load. These complementary models help distinguish pressure-dependent from AngII-mediated vascular mechanisms and provide a platform to develop targeted therapies for coronary vasculopathy and AngII-driven vascular disease.

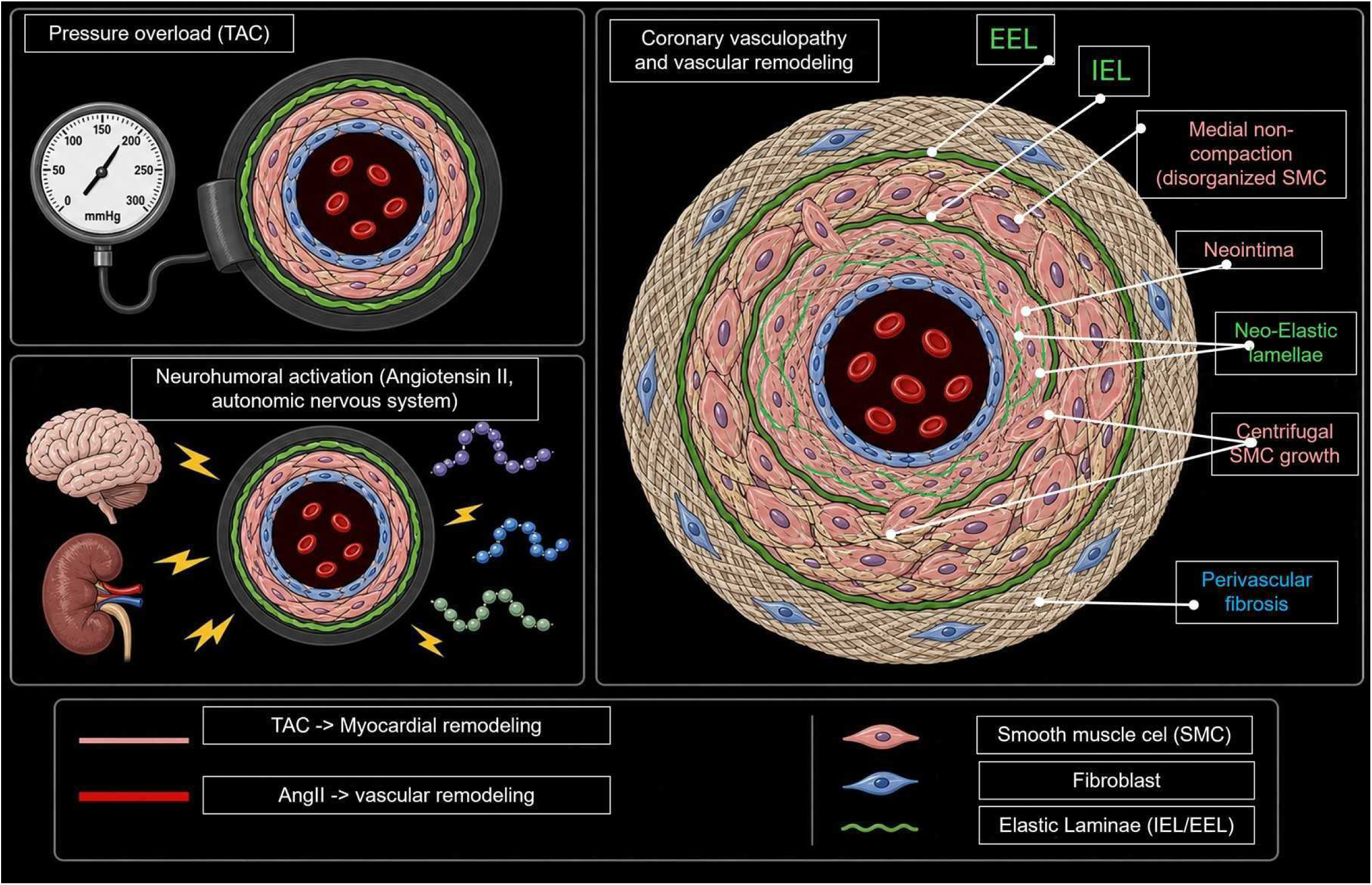

## Introduction

### Cardiac Remodeling: Pathophysiological Overview

Pathological cardiovascular remodeling encompasses structural and functional alterations in both the myocardium and the coronary vasculature, typically progressing from inflammation to fibrosis. Key features include cardiomyocyte hypertrophy, fibroblast activation, excessive extracellular matrix (ECM) deposition, and parallel changes in intramyocardial vessels that together stiffen the ventricular wall and impair coronary flow reserve [1, 2]. These processes are highly relevant to clinical syndromes such as heart failure with preserved ejection fraction (HFpEF), where myocardial and vascular remodeling coexist and contribute to diastolic dysfunction [3–6].

### Hemodynamic versus neurohumoral drivers have distinct vascular impact

In the absence of major metabolic or genetic disease, two broad classes of stimuli dominate vascular remodeling process: hemodynamic overload and neurohumoral activation. Chronic pressure overload, such as long-standing non–RAAS–driven hypertension, severe aortic stenosis, or coarctation, imposes increased wall stress, leading to stretch–strain injury, cardiomyocyte hypertrophy, and interstitial fibrosis [7–13]. In parallel, activation of the renin–angiotensin–aldosterone system (RAAS) represents a systemic neurohumoral stimulus that disproportionately targets the vasculature. Angiotensin II (Ang II), the principal RAAS effector, not only promotes hypertrophy and fibrosis through profibrotic signaling in interstitial cells [11, 14–16] but also directly modulates vascular smooth muscle cells (VSMCs), endothelial cells (EC), and adventitial fibroblasts. Clinically, chronic Ang II elevation is a hallmark of conditions such as chronic kidney disease and renovascular hypertension, where coronary and systemic small-artery disease, neointima formation, and elastin injury frequently accompany myocardial remodeling. Current vasculopathy therapies primarily target VSMC tone (e.g., vasodilators, antispasmodics), but largely neglect VSMC proliferation, phenotype plasticity, and intercellular communication—central drivers of vascular stiffness, neointima formation, and impaired coronary flow that are increasingly recognized across atherosclerotic and non-atherosclerotic vasculopathies.

### Myocardial vs. Vascular Remodeling

Research has disproportionately emphasized myocardial fibrosis and stiffness while underappreciating coronary vasculopathy and perivascular fibrosis, which are frequently conflated with “myocardial” pathology [4, 13, 14]. Yet these represent distinct cardiac compartments. The myocardium comprises cardiomyocytes, interstitial fibroblasts, and capillaries (with pericytes, EC, and VSMCs), whereas the coronary vasculature encompasses EC, medial VSMCs, adventitial fibroblasts, internal and external elastic laminae (IEL, EEL), and perivascular ECM [15, 16]. Clinical and experimental data suggest that pressure-dominant stimuli preferentially drive diffuse myocardial hypertrophy and interstitial fibrosis, whereas AngII–dominant states are particularly prone to small-artery remodeling, neointima formation, and elastin injury, even when occurring at more modest pressure loads [17–19]. The initiating events that drive vascular remodeling remain controversial. Classical atherosclerosis emphasizes endothelial injury and lipid-rich plaque formation as triggers of centripetal neointima growth and luminal narrowing. In AngII–rich states, however, VSMCs and adventitial cells are exposed to high circulating and tissue-level AngII, raising the possibility that vasculopathy can arise primarily from VSMC-centric processes beneath a structurally preserved endothelium. Although VSMCs were traditionally regarded as passive repair mediators, emerging evidence positions them as active, drivers of vascular disease, capable of transitioning into mesenchymal, inflammatory, or macrophage-like states that remodel the intima, media, and elastic laminae. Our experimental models isolate these pathologic processes to identify vasculopathy-specific therapeutic vulnerabilities [8, 20]. The directionality of vascular remodeling—centripetal (intima → vessel lumen) versus centrifugal (intima→ adventitia)—remains incompletely defined across different vasculopathies [21–23]. Canonical stimuli include inflammation, thrombosis, and mechanical injury, which trigger endothelial activation or denudation via reactive oxygen species and biomechanical stress. In atherosclerosis, this paradigm centers on endothelial dysfunction and plaque formation; in AngII–driven vasculopathy, the balance may shift toward VSMC-mediated neointima formation, elastic lamina remodeling, and perivascular fibrosis, even in the absence of overt endothelial loss [24, 25].

##### Manuscript Highlights

###### Key Findings

- Hemodynamic and neurohumoral stimuli distinctly drive myocardial versus coronary remodeling
- Coronary vasculopathy comprises neointima formation, medial VSMC changes, elastic lamina disruption, inflammation, and perivascular fibrosis
- Myocardial and vascular remodeling pathways diverge mechanistically and temporally
- Current vasculopathy therapies target VSMC relaxation but neglect proliferation, phenotype switching, and intercellular communication

###### Key Definitions

- **Cardiac remodeling**: Integrated myocardial and vascular structural/functional adaptations
- **Myocardial remodeling**: Cardiomyocyte hypertrophy/necrosis (or apoptosis?) plus interstitial ECM deposition by fibroblasts/inflammatory cells
- **Coronary remodeling**: Multilayer vascular changes spanning the intima, tunica media (SMC-driven), adventitia, elastic laminae (IEL/EEL), and perivascular matrix
- **Hemodynamic stress**: Pressure/strain-induced biomechanical cardiomyocyte and vascular injury
- **Neurohumoral stress**: Direct cellular effects of hormones (AngII) independent of mechanical load

### Study Objectives

This study was designed to dissect the independent and synergistic contributions of hemodynamic overload and AngII–dominant neurohumoral stress to myocardial versus coronary remodeling, with particular emphasis on VSMC-centric vasculopathy. We used murine models that preferentially accentuate pressure-overload myocardial remodeling (transverse aortic constriction, TAC) or AngII–dominant coronary vasculopathy (AngII/phenylephrine and high-dose AngII), mirroring clinical contexts of fixed afterload and chronic RAAS activation, respectively, while acknowledging that each stimulus engages overlapping pathways. By combining invasive hemodynamics with multiregion histologic and lineage-tracing analyses, we mapped early inflammatory events, VSMC phenotypic transitions, neointima and elastic lamina remodeling, and late fibrosis within distinct cardiac compartments. Rather than providing a comprehensive model of HFpEF, this framework identifies AngII–associated vascular remodeling programs that are relevant to HFpEF-relevant, AngII–elevated disease states such as chronic kidney disease and renovascular hypertension and offers experimental platforms to explore therapies that specifically target coronary vasculopathy and perivascular fibrosis alongside classic pressure-overload cardiomyopathy.

## Results

### 1. Hemodynamic load and global cardiac remodeling in TAC versus AngII models

***<u>1.1</u>*** Hemodynamic responses to pressure overload and AngII-dominant stress Mice underwent TAC surgery, AngII/PE infusion (1.5 µg/g/day, 50 μg/g/day), or HD-AngII (3 µg/g/day) to induce neurohumoral or hemodynamic stress (**Fig. 2A–C**). TAC markedly elevated LV systolic pressure compared to untreated mice, while AngII/PE and HD-AngII produced more moderate pressure increases that scaled inversely with AngII dose. No significant sex-related changes in hemodynamics were observed (blue symbols, males; red symbols, females). All stress models elevated LV end diastolic pressure (LVEDP) above control levels, although no differences between groups were observed. Right ventricular systolic pressures demonstrated near-normal values (28 ± 2 vs 20 ± 4 mmHg in controls), measured but not shown.

**Figure 1.**
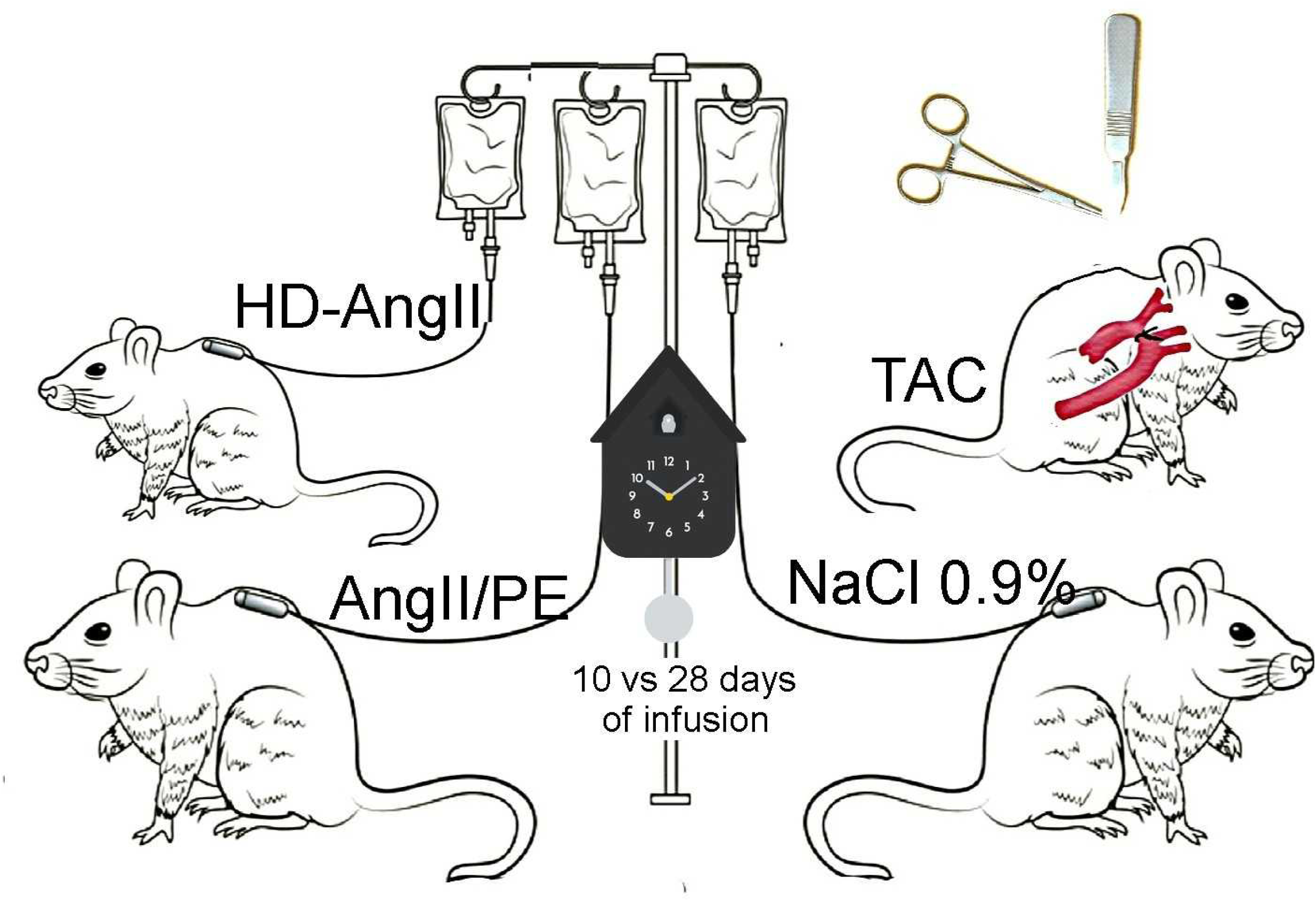
Mouse models of cardiac remodeling. Transverse aortic constriction (TAC; 28 days) induces pressure overload–mediated remodeling. AngII (1.5 µg/g/day) plus phenylephrine (50 µg/g/day) produces an HFpEF model, and high-dose AngII (HD-AngII; 3 µg/g/day for 10 or 28 days) generates a vasculopathy-predominant remodeling phenotype.

**Figure 2.**
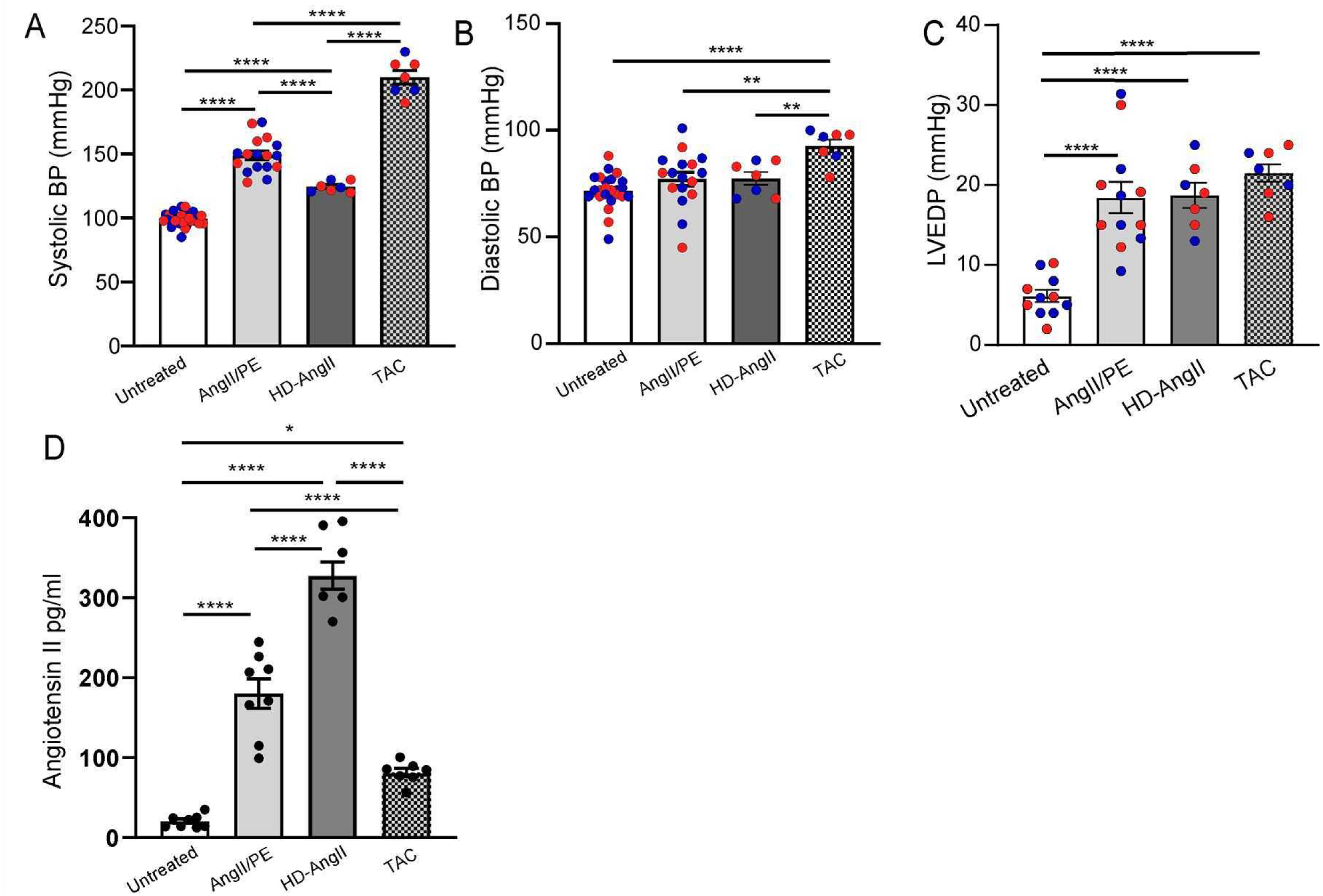
Hemodynamic assessment across cardiac remodeling models. A-C) Retrograde left ventricular (LV) catheterization-derived systolic pressure and LV end-diastolic pressure (LVEDP) in TAC (after 28 days), AngII/PE and HD-AngII (after 28 days) groups. (A) TAC surgery produced the highest systolic and (B) diastolic blood pressure. (C) LVEDP remained comparable across all experimental conditions. Group sizes were n≥7 per mouse model. Statistical significance was defined as: * P<0.05; ** P<0.01, and *** P<0.001. Data was analyzed using two-way ANOVA with multiple comparisons. No significant sex-related differences (males are shown in blue and females in red) in hemodynamic parameters were observed in either untreated or treated groups across all models.

#### Circulating Ang II levels across TAC and Ang II-infusion models

To quantify neurohumoral activation in each model, we measured circulating AngII concentrations in serum from control, TAC, AngII/PE, and HD-AngII mice at day 28. Ang II levels were low in control mice and significantly elevated in all three stress models, with the highest concentrations observed in HD-AngII, intermediate levels in AngII/PE, and modest but still elevated levels in TAC (**Fig. 2D**).

### 1.2 Effects of TAC and AngII on myocardial hypertrophy

Increased cardiomyocyte cross-sectional area (**Fig. 3A-C**) and heart weight-to-tibia length ratio (**Fig. 3D**) confirmed cardiomyocyte hypertrophy across TAC, AngII/PE, and HD-AngII groups. TAC induced uniform hypertrophy throughout the LV free wall and septum, whereas AngII/PE and HD-AngII produced patchy cardiomyocyte hypertrophy primarily co-localized to fibrotic regions. TAC also elicited greater cardiomyocyte hypertrophy compared to the AngII groups (**Fig. 3A-C**). We initially tested phenylephrine as an isolated hemodynamic stressor using continuous α1-adrenergic stimulation. This approach produced marked systolic hypertension (to ∼140 mmHg) but failed to induce significant myocardial or vascular remodeling (**Supplemental figure 1**), even after a fourfold dose escalation. At doses above 100 µg/kg/day, pressor responses waned, likely due to local ischemia and poor drug absorption. In contrast to a recent bolus phenylephrine study reporting rapid fibrosis and increased EF under nonhypertensive conditions [26], our findings with continuous infusion were inconsistent and difficult to interpret. We therefore adopted TAC as a more robust and reproducible model of hemodynamic cardiac stress for the present study.

**Figure 3.**
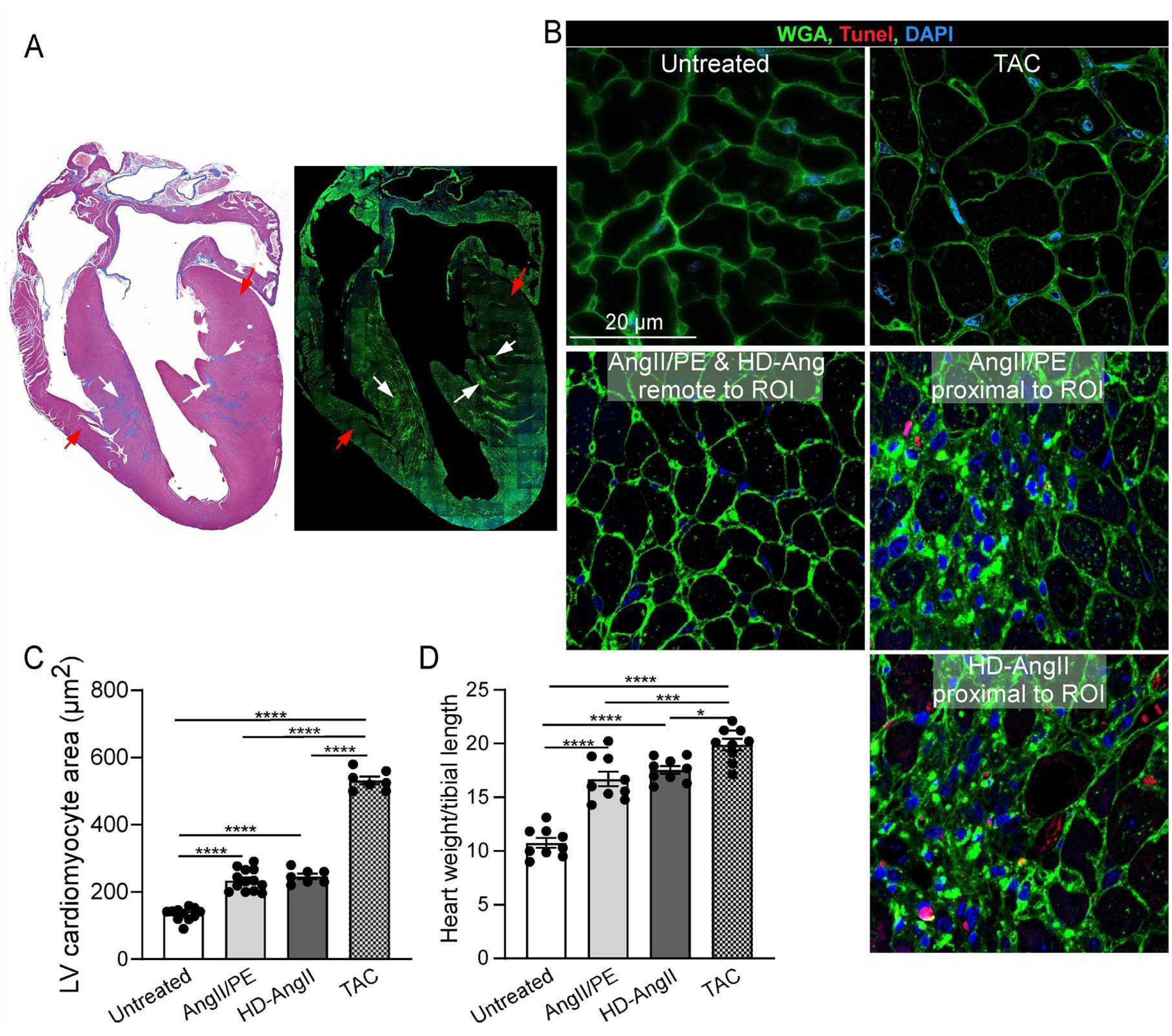
Cardiomyocyte hypertrophy and death across remodeling models. (A) Low-magnification Masson’s trichrome and IF images of the whole heart provide an anatomic roadmap for the higher-magnification insets shown in subsequent panels (white arrows – area proximal to ROI; red arrows - area remote to ROI). (B) Representative wheat germ agglutinin (WGA, green), TUNEL (red), and DAPI (blue) staining of left ventricular cardiomyocytes 28 days after AngII/PE or HD-AngII infusion (regions adjacent to and remote from fibrosis/inflammation) or 28 days after TAC surgery. AngII/PE and HD-AngII hearts exhibit heterogeneous cardiomyocyte hypertrophy, with marked CM enlargement, CM necrosis in ROIs and near-normal CM size in remote zones. (C) Quantification of myocyte cross-sectional area adjacent to fibrotic regions showed increased hypertrophy in AngII/PE and HD-AngII groups at day 28; TAC demonstrates more homogeneous enlargement throughout the myocardium. (D) Heart weight-to-tibial length ratios are elevated across all treatment groups, with TAC showing the highest values. Group sizes were n≥7 per mouse model. Statistical significance was defined as: * P<0.05; ** P<0.01, and *** P<0.001. Data was analyzed using two-way ANOVA with multiple comparisons.

### 1.3 Effects of TAC and AngII on cardiac fibrosis patterns

Cardiac fibrosis manifested as predominant interstitial, perivascular, or mixed phenotypes (**Fig. 4A1-4).** Qualitatively, AngII/PE showed interstitial (∼55% of mice) (**Fig. 4B1**) [27], VSMC hypertrophy and perivascular fibrosis (∼30% of mice) and mixed (∼15% of mice) phenotypes, while HD-AngII was dominated by VSMC hypertrophy and perivascular fibrosis (∼80% of mice) (**Fig. 4B2**). TAC induced predominantly a mild interstitial fibrosis phenotype (∼70% of mice), with minimal perivascular involvement (**Fig. 4B3**). Quantitative analysis confirmed that the highest interstitial and perivascular fibrosis burden was in HD-AngII, followed by AngII/PE, with TAC exhibiting the least fibrosis overall (**Fig. 4C, D**).

**Figure 4.**
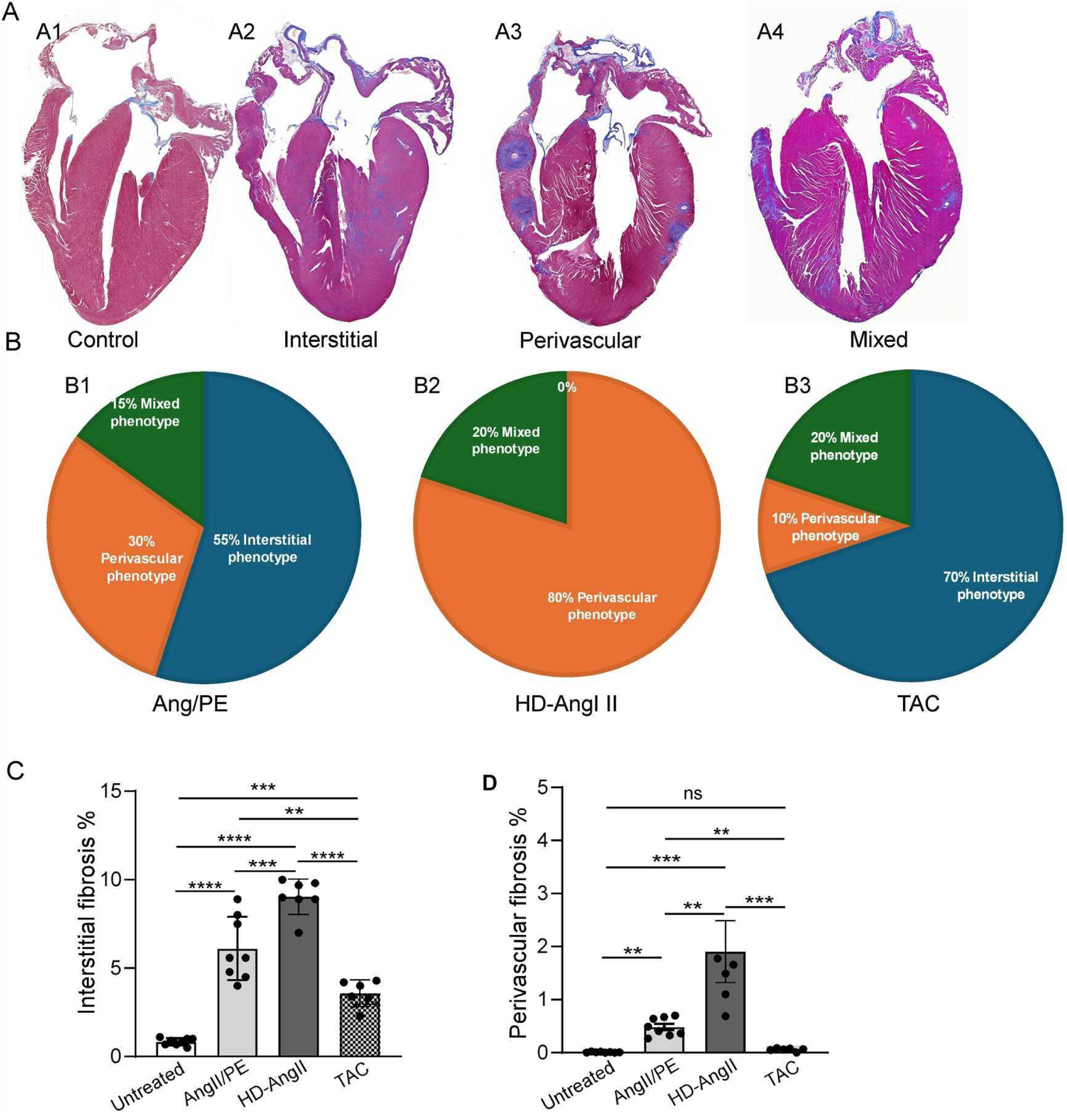
Cardiac fibrosis phenotypes across experimental models. (A) Representative Masson’s trichrome-stained myocardial sections demonstrate interstitial, perivascular, and mixed fibrosis phenotypes following 28 days of AngII/PE infusion, 28 days after TAC surgery, or 28 days of HD-AngII infusion. (B) Distribution of fibrosis phenotypes reveals perivascular predominance in the HD-AngII model and interstitial predominance in TAC. (C) Quantitative analysis of interstitial and perivascular fibrosis (ECM relative to total coronary section area) shows the greatest overall fibrosis in HD-AngII models across both compartments, with TAC exhibiting minimal perivascular involvement. Group sizes were n≥6 per each mouse model and cardiac remodeling phenotype. Statistical significance was defined as: * P<0.05; ** P<0.01, and *** P<0.001. Data was analyzed using two-way ANOVA with multiple comparisons.

## 2. AngII-dominant coronary vasculopathy and perivascular fibrosis

### 2.1 Temporal progression of coronary vascular remodeling and inflammation

As previously described, myocardial interstitial changes in AngII/PE-treated mice began with inflammation from days 3–10, transitioning to fibrosis by day 28 [27]. The prominent vascular phenotype in HD-AngII mice at day 28 prompted examination at an earlier time point (**Fig. 5A, A’**). By 10 days, perivascular regions showed early inflammatory patterns with macrophage infiltration, persisting into later stages alongside the progressing fibrotic changes (**Fig. 5B-D**). Inflammatory severity and macrophage activation were quantified using CD68 staining intensity in regions of interest (**Fig. 5E**), confirming significant infiltration at both early and late AngII stages. Extracellular perivascular matrix alterations consistently accompanied changes in VSMC proliferation and hypertrophy.

**Figure 5.**
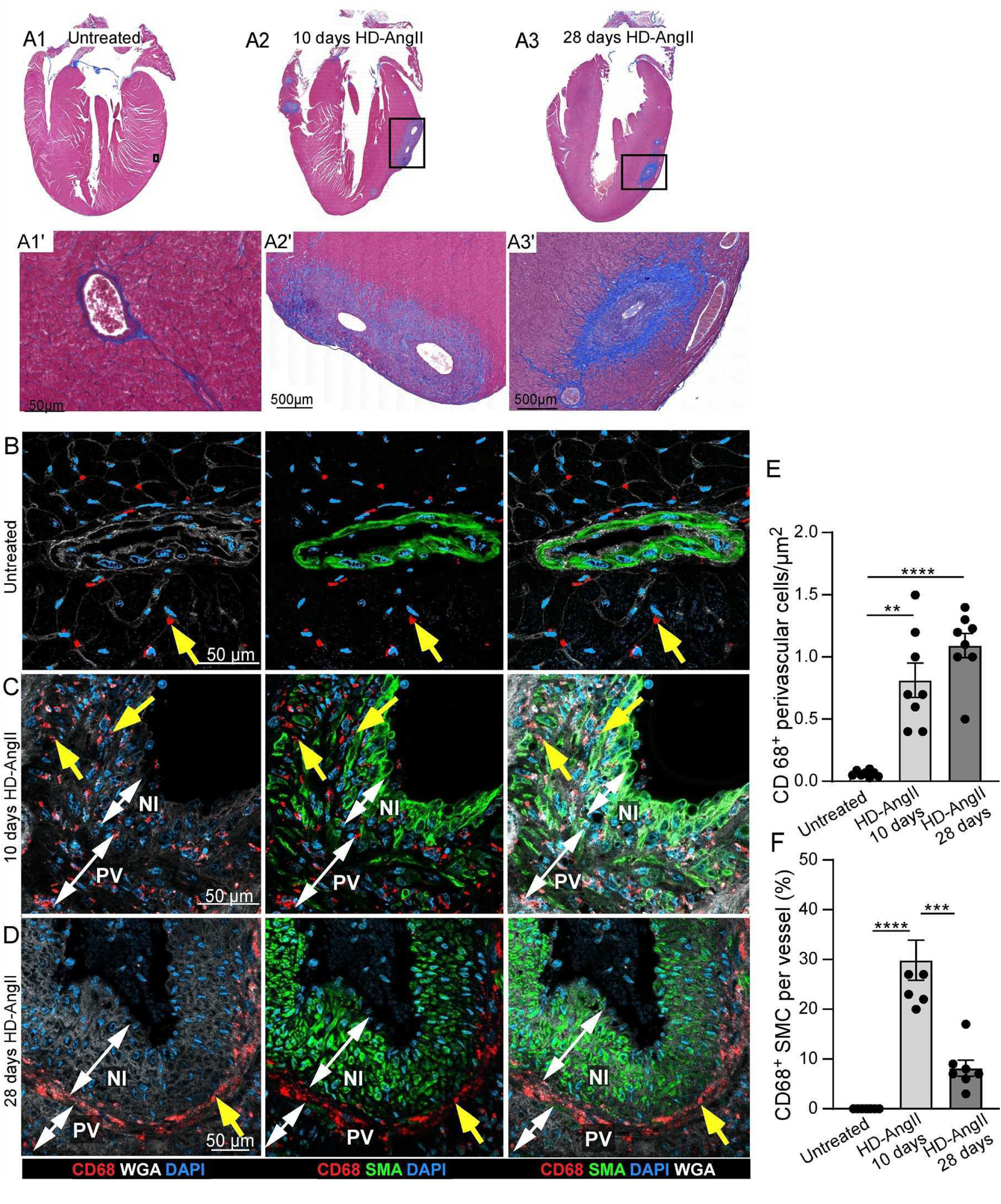
Temporal progression of perivascular inflammation, fibrosis, and SMC-to-macrophage phenotypic transition during HD-AngII infusion. (A) Representative Masson’s trichrome staining at 10 and 28 days reveals progressive extracellular matrix deposition and cellular infiltrates in perivascular regions (A1’-3’: inset magnifications). (B–D) Immunofluorescence for CD68 (red; perivascular macrophages/CD68+ +SMCs, yellow arrows), wheat germ agglutinin (WGA, white), smooth muscle actin (SMA, green), and DAPI (blue) shows persistent perivascular macrophage accumulation from early (day 10) through late (day 28) stages, with maturation of perivascular fibrosis over time; white double arrows define neointima (NI) and perivascular (PV) areas. (E) Quantitative analysis confirms sustained macrophage infiltration (per µm^2^) across both early and late phases of HD-AngII infusion. (F) Quantification reveals approximately one-third of SMCs acquire macrophage markers by day 10, with the process declining at later time points by day 28. Group sizes were n≥8 at 10 and 28 days of HD-AngII treatment. Statistical significance was defined as: * P<0.05; ** P<0.01, and *** P<0.001. Data was analyzed using two-way ANOVA with multiple comparisons.

### 2.2 VSMC phenotypic transitions in AngII-induced vasculopathy

Several studies examining VSMC phenotypes in atherosclerosis have demonstrated transitions from a mature contractile phenotype to alternative states [28]. To determine whether coronary artery VSMCs undergo similar transitions in response to AngII, we performed immunostaining for CD68 to identify macrophage-like phenotypes and SCA1 and KLF4 to identify undifferentiated/mesenchymal phenotypes. Expression of macrophage markers in VSMC coincided with perivascular macrophage infiltration on day 10 of HD-AngII infusion (**Fig. 5B, C**). However, with prolonged AngII exposure, macrophage marker expression within VSMCs declined despite persistent perivascular macrophage accumulation (**Fig. 5B, D**). Quantitative analysis showed that CD68^+^ VSMC transformation peaked during early AngII infusion and decreased by day 28 (**Fig. 5F**). Increased expression of mesenchymal markers, KLF4 and SCA1, were detectable by day 7 and persisted through day 28 (**Fig. 6A–C**). Quantitative analysis indicated that the undifferentiated, KLF4+, SCA1+, phenotype peaked during early AngII infusion, suggesting a transient stem-like state during active vascular remodeling (**Fig. 6D, E**). To assess changes in proliferation and quiescence in response to HD-AngII infusion, sections were immunostained for Ki67 and CDKN1A (p27^Kip1^). In control mice, coronary vessels did not express Ki67 and expressed high levels of CDKN1A (**Fig. 7A1, B, E, F**). After 10 days HD-AngII infusion, coronary vessels expressed high levels of Ki67 and very low levels of CDKN1A (**Fig. 7A2, C, E, F**). After 28 days HD-AngII infusion, Ki67 expression returned to near baseline levels and CDKN1A remained at low levels (**Fig. 7A3, D, E, F**).

**Figure 6.**
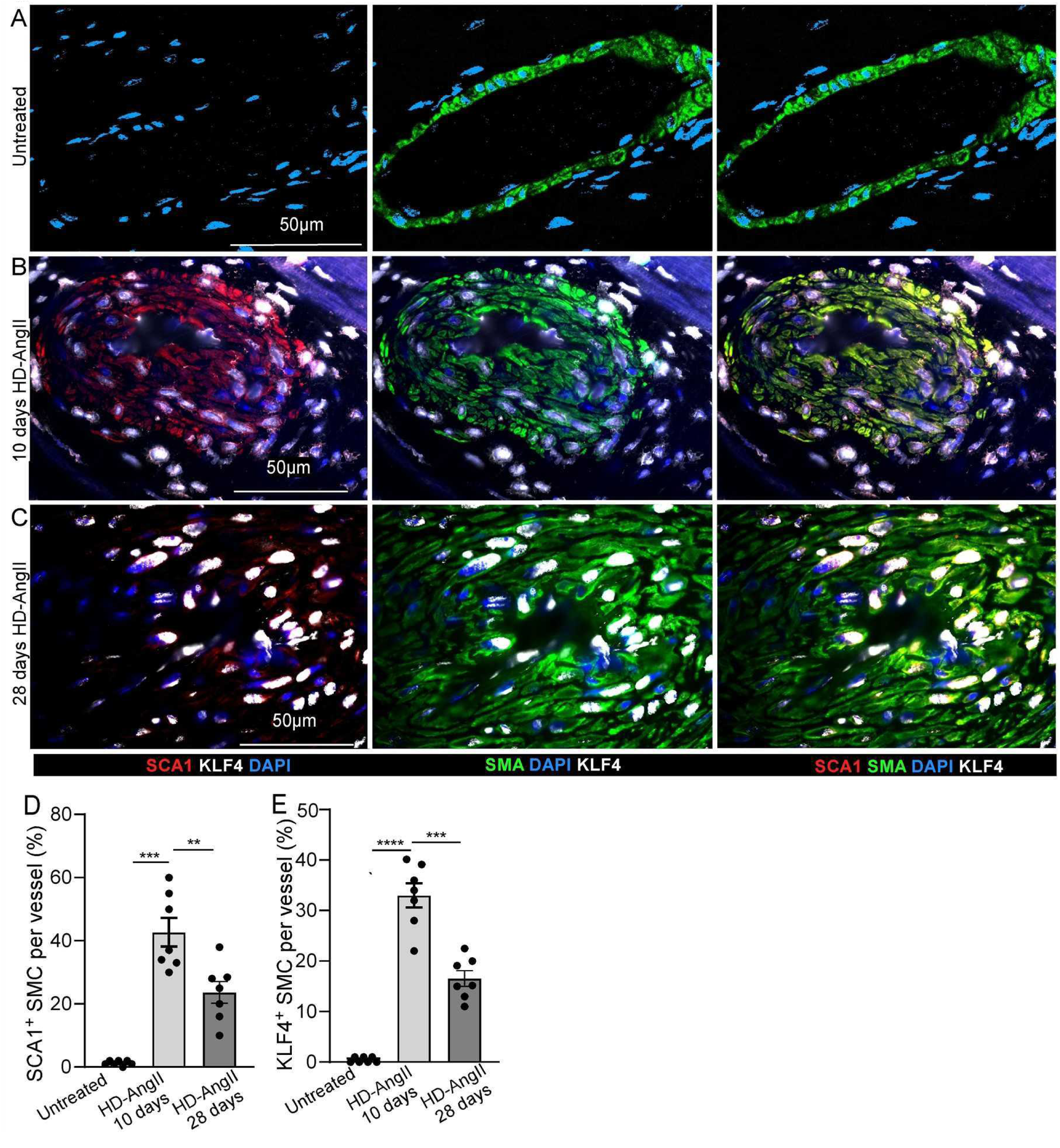
VSMC dedifferentiation in response to AngII infusion. A-C) Immunofluoresce staining for smooth muscle actin (SMA; green), SCA1 (red, red arrows), KLF4 (white, white arrows), and DAPI (blue) reveals SMC transition from differentiated to undifferentiated phenotype, marked by co-expression of SCA1 and KLF4. Individual channel views showing KLF4/SMA/DAPI, SCA1/KLF4/DAPI, and SCA1/KLF4/SMA/DAPI staining patterns. D) Quantification of SCA1+ and SMA+ cells per vessel. E) Quantification of KLF4+, SMA+ cells per vessel. Group sizes were n≥8 at 10 and 28 days of HD-AngII treatment. Statistical significance was defined as: * P<0.05; ** P<0.01, and *** P<0.001. Data was analyzed using two-way ANOVA with multiple comparisons.

**Figure 7.**
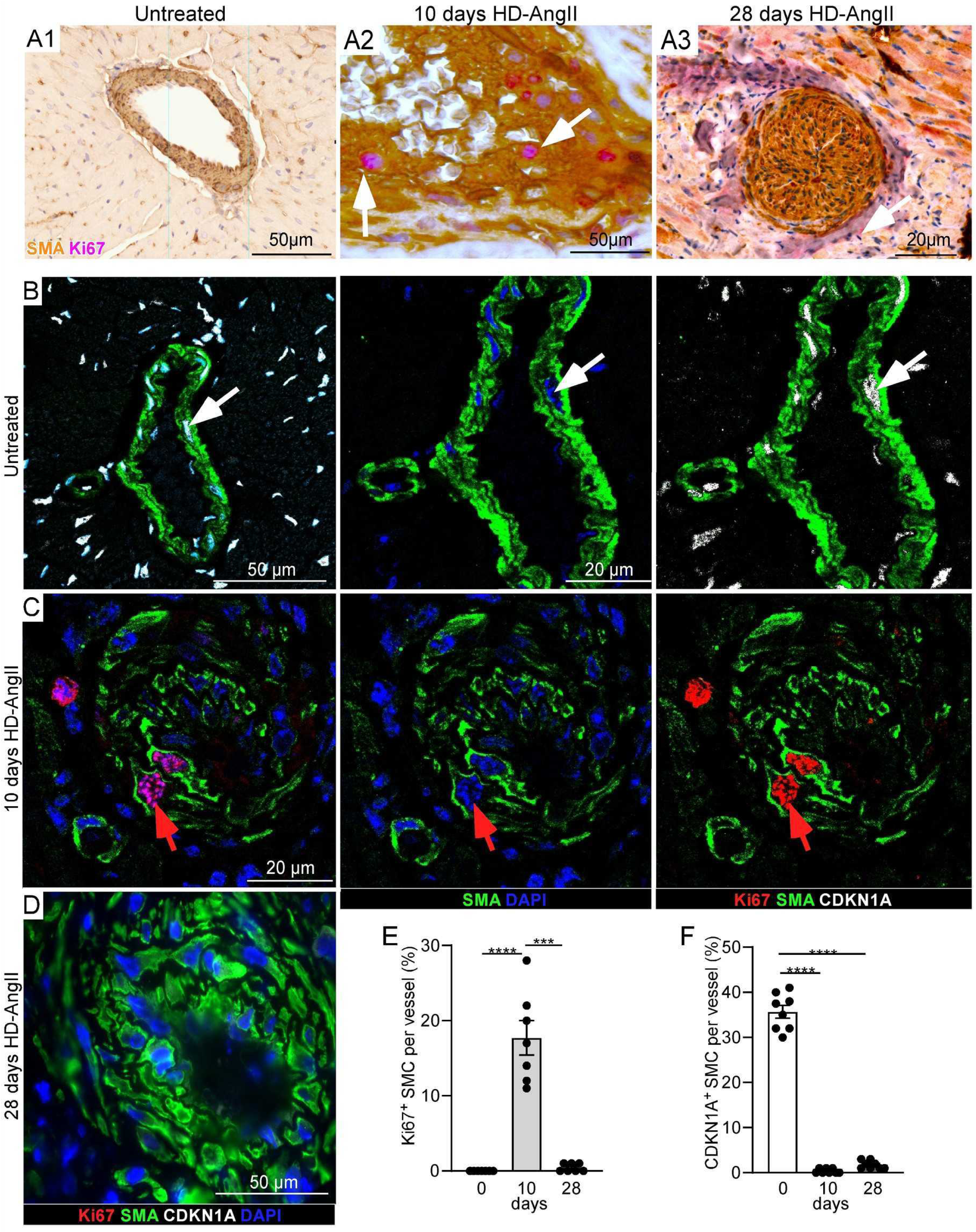
SMC proliferation and quiescence dynamics during HD-AngII infusion. (A) Bright-field immunohistochemistry for Ki67 (pink, white arrows) and SMA (brown) reveals SMC proliferation at early stages. (B-D) Immunofluorescence for Ki67 (red, red arrows) and CDKN1a (white, white arrows) shows senescent SMCs (CDKN1a+ nuclei) in controls, proliferative states (Ki67+ nuclei) at early infusion (7–10 days), and absence of proliferation at late stages. Individual channel views of CDKN1a/SMA, SMA/DAPI, Ki67/SMA, and SMA/DAPI staining. (E) Quantification show peak of proliferative state (Ki67+, SMA+ cells per vessel) at 10 days of HD-AngII infusion. (F) Quantification showing peak quiescence (CDKN1A+, SMA+ cells per vessel) of VSMC in untreated mice. Group sizes were n≥8 at 10 and 28 days of HD-AngII treatment. Statistical significance was defined as: * P<0.05; ** P<0.01, and *** P<0.001. Data was analyzed using two-way ANOVA with multiple comparisons.

### 2.3 Neointima formation, elastic lamina disruption, and vascular structural changes in the absence of overt endothelial layer alterations

***2.3A*** To determine whether EC contribute directly to neointima formation in AngII-driven coronary vasculopathy, we crossed the Cdh5-CreERT2 (VE-cadherin) [29] endothelial targeting allele mice with the *ROSA^tdTomato^*(Ai9) [30] reporter allele and analyzed coronary arteries for tdTomato expression in animals with advanced vascular lesions. In Cdh5-CreERT2; *ROSA^tdTomato^* mice given tamoxifen chow (TAM) from 3-5 weeks of age and induced with AngII at 8 weeks of age (**Fig. 8A**). After 10 or 28 days, coronary vessels were imaged for tdTomato, smooth muscle actin (SMA) and elastin. ECs showed robust tdTomato labeling along an intact luminal monolayer in both control and AngII-treated hearts (**Fig. 8B**). Despite severe concentric neointimal thickening in affected vessels, we did not detect tdTomato-positive cells within the neointima, indicating an absence of endothelial-derived cells in the lesion core (**Fig. 8B**). The endothelial layer overlying neointimal lesions remained continuous without evidence of rupture, denudation, or large gaps by histologic and fluorescence microscopy assessment.

**Figure 8.**
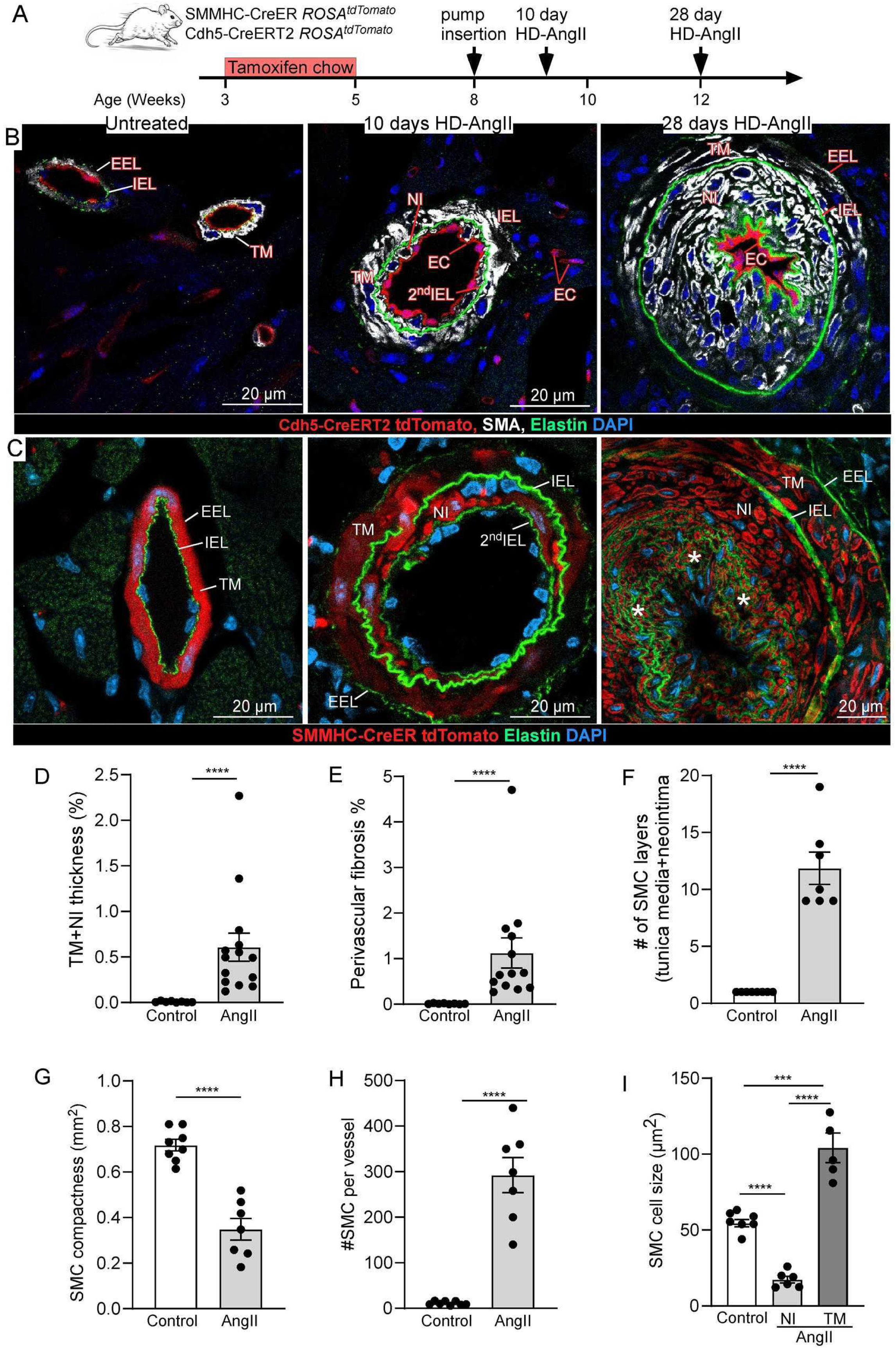
Vascular elastic membrane remodeling and endothelial/smooth muscle cell dynamics during HD-AngII infusion. (A) Experimental plan for SMC and EC lineage–tracing, showing SMMHC-CreER and Cdh5-CreERT2;ROSA^tdTomato^ induction with tamoxifen at 3–5 weeks of age, followed by a 4-week washout period before HD-AngII minipump implantation. (B) Endothelial lineage tracing in AngII-induced coronary vasculopathy. Endothelial cell dynamics were assessed using Cdh5-CreERT2; ROSA^tdTomato^ mice (EC lineage label, red) in relation to the elastic lamina (elastin, green) at early (7–10 days) and late (28 days) stages of HD-AngII infusion. (C) SMC lineage labeling (ROSA-tdTomato, red) in AngII-induced coronary vasculopathy. SMC dynamics were assessed using SMMHC-CreER; *ROSA^tdTomato^*(red), and elastin (green). (D–I) Quantitative morphometry of immunofluorescence images in HD-AngII–treated mice: (D) HD-AngII infusion induced marked concentric remodeling, with a significant increase in combined tunica media plus neointima thickness compared with controls; (E) perivascular fibrosis was substantially higher in AngII-treated vessels; (F) the number of SMC layers across the tunica media and neointima was markedly increased following AngII infusion; (G) AngII treatment reduced SMC compactness, consistent with a more disorganized medial architecture; (H) total SMC number per vessel was significantly elevated in treated arteries; and (I) SMC cross-sectional area was also increased, with hypertrophy evident in both neointimal and medial SMCs relative to untreated controls. Group sizes were n≥6 at 10 and 28 days of HD-AngII treatment. Statistical significance was defined as: * P<0.05; ** P<0.01, and *** P<0.001. Data was analyzed using two-way ANOVA with multiple comparisons.

***2.3B*** To identify the fate of VSMC in response to HD-AngII infusion, SMMHC-CreER; *ROSA^tdTomato^* (VSMC-lineage) lineage-trace mice [30] were given TAM chow from 3-5 weeks of age and induced with AngII at 8 weeks of age (**Fig. 8A**). After 10- or 28-days coronary vessels were imaged for tdTomato and elastin. Coronary vessels in untreated control mice showed a tdTomato+ tunica media (TM) just outside the internal elastic lamina (IEL) (**Fig. 8C**). After 10 days HD-AngII infusion, tdTomato+ cells were observed internal to the IEL and a secondary weakly stained IEL formed (2^nd^IEL) (**Fig. 8C**). At the 10-day time point HD-AngII infusion markedly disrupted coronary artery extracellular matrix organization, including deformation and rupture of both the internal IEL) and secondary internal (2^nd^IEL) elastic laminae (**Fig. 8C; Supplemental Videos 1–2**). These data suggest that VSMCs proliferated centripetally through breaks in the IEL, forming non-compact, multilayered cell strata interspersed with disorganized secondary-elastic fibers forming a neo-intima (NI). After 28 days HD-AngII infusion, the NI was markedly expanded with tdTomato+ cells, accompanied by outward displacement of the elastic laminae, thickening of the TM, and prominent perivascular fibrosis (**Fig. 5A3, 8C**) Quantitative analyses confirmed pronounced changes in VSMC morphology and organization (**Fig. 8D-I)**, smaller (hypotrophic) VSMCs in the neointima and larger (hypertrophic) VSMCs in the medial layer (**Fig. 8I**), and increased perivascular fibrosis (**Fig. 8E**). Importantly, myocardial ECM remodeling at both early and late HD-AngII time points did not involve new elastic fiber deposition (**Supplemental Fig. 2**).

### 2.4 Elastin breakdown and desmosine as markers of AngII-driven vasculopathy

To assess remodeling of tissue elastin at 10 days of HD-AngII infusion, we measured serum desmosine as an index of elastin degradation [31]. Compared to controls, mice infused with HD-AngII for 28 days showed increased serum desmosine levels (**Fig. 9A**). Stratification of HD-AngII infused mice by histologic phenotype (vascular vs interstitial) showed that desmosine levels were significantly higher in HD-AngII mice with a predominant vascular phenotype compared to HD-AngII mice exhibiting an interstitial phenotype (**Fig. 9A**). These findings indicate greater elastin breakdown in animals developing early perivascular/vascular remodeling than in those with primarily interstitial myocardial fibrosis. Consistent with these observations, analysis of urinary elastin degradation products demonstrated elevated desmosine levels in HD-AngII–treated animals; however, no significant differences were observed between phenotypic subgroups (**Fig. 9B**).

**Figure 9.**
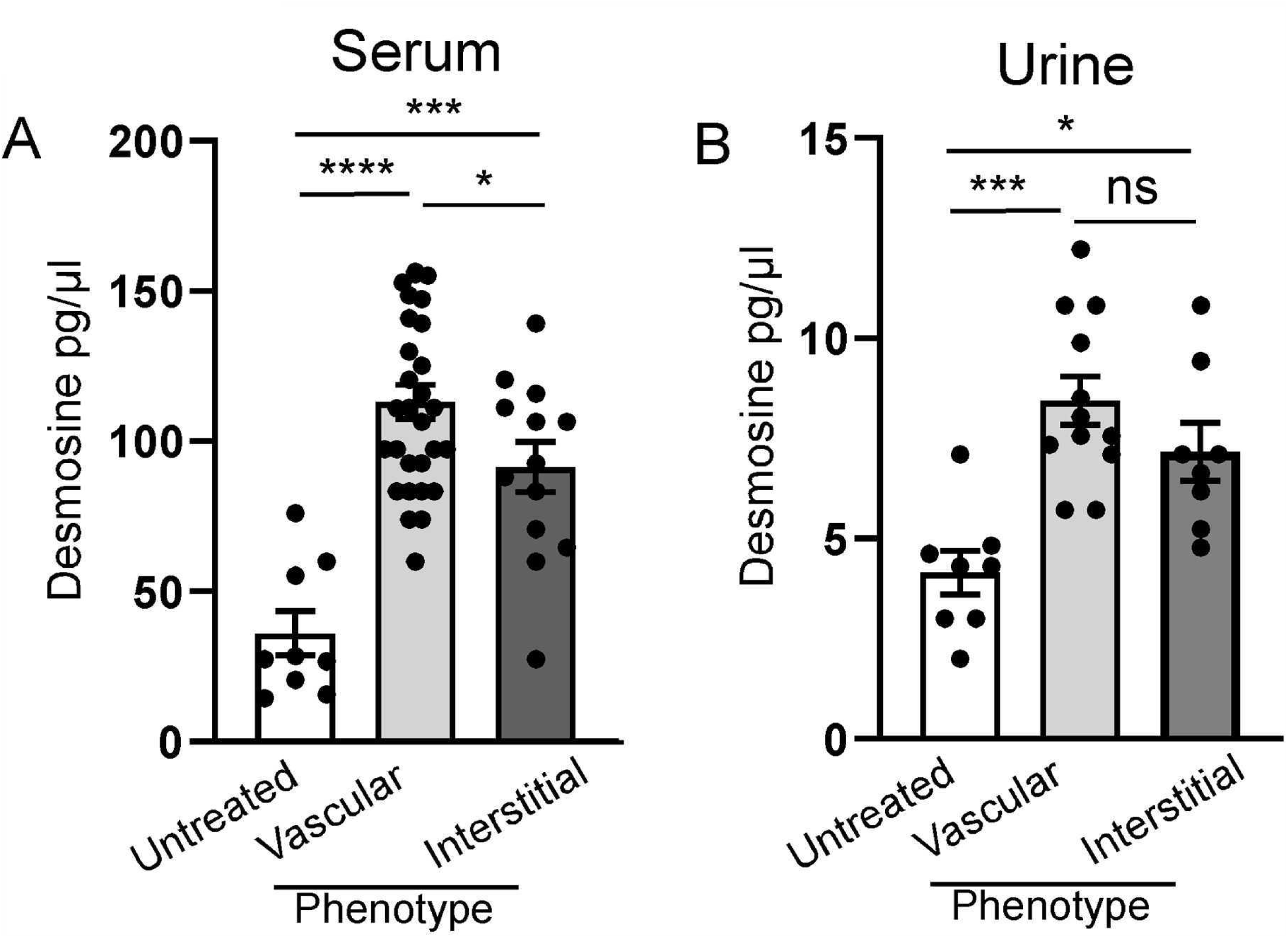
Serum and urine desmosine levels across cardiac remodeling phenotypes. (A) After 28 days of HD-AngII infusion, serum desmosine concentrations were highest in mice with a vascular phenotype and showed only a modest increase in those with an interstitial phenotype compared with untreated controls (N=9 untreated, N=28 vascular, N=13 interstitial). (B) After 28 days of HD-AngII infusion, urine desmosine concentrations were similarly elevated in treated mice irrespective of histologic subtype, with no significant differences between vascular and interstitial phenotypes (N=8 untreated, N=12 vascular, N=8 interstitial). Statistical significance was defined as: * P<0.05; ** P<0.01, and *** P<0.001. Data was analyzed using two-way ANOVA with multiple comparisons.

### 2.5 AT1R–AKT signaling in VSMCs during AngII-induced vasculopathy

AT1R, which is essential for cardiac development and function [32, 33], was detectable by immunostaining in cardiomyocytes (LV and LA), vascular smooth muscle cells, ECs, and inflammatory cells in both control and AngII-treated hearts (**Fig. 10A, B**). These staining patterns were interpreted qualitatively and were used primarily to illustrate the broad cardiovascular distribution of AT1R rather than to quantify differences in expression between groups. Downstream pathway analysis focusing on MAPK and AKT signaling showed robust AngII-induced activation of AKT in VSMCs. Phospho-AKT staining was minimal in control vessels but markedly increased in AngII-infused mice, localizing mainly to the plasma membrane and cytoplasm of VSMCs (**Fig. 10C-E**). This activation, consistent with prior reports in AngII-stimulated vasculature [34–36], emerged by day 10 and persisted through day 28, with similar magnitude and distribution with short- and long-term infusion, indicating that AKT activation was independent of infusion duration.

**Figure 10.**
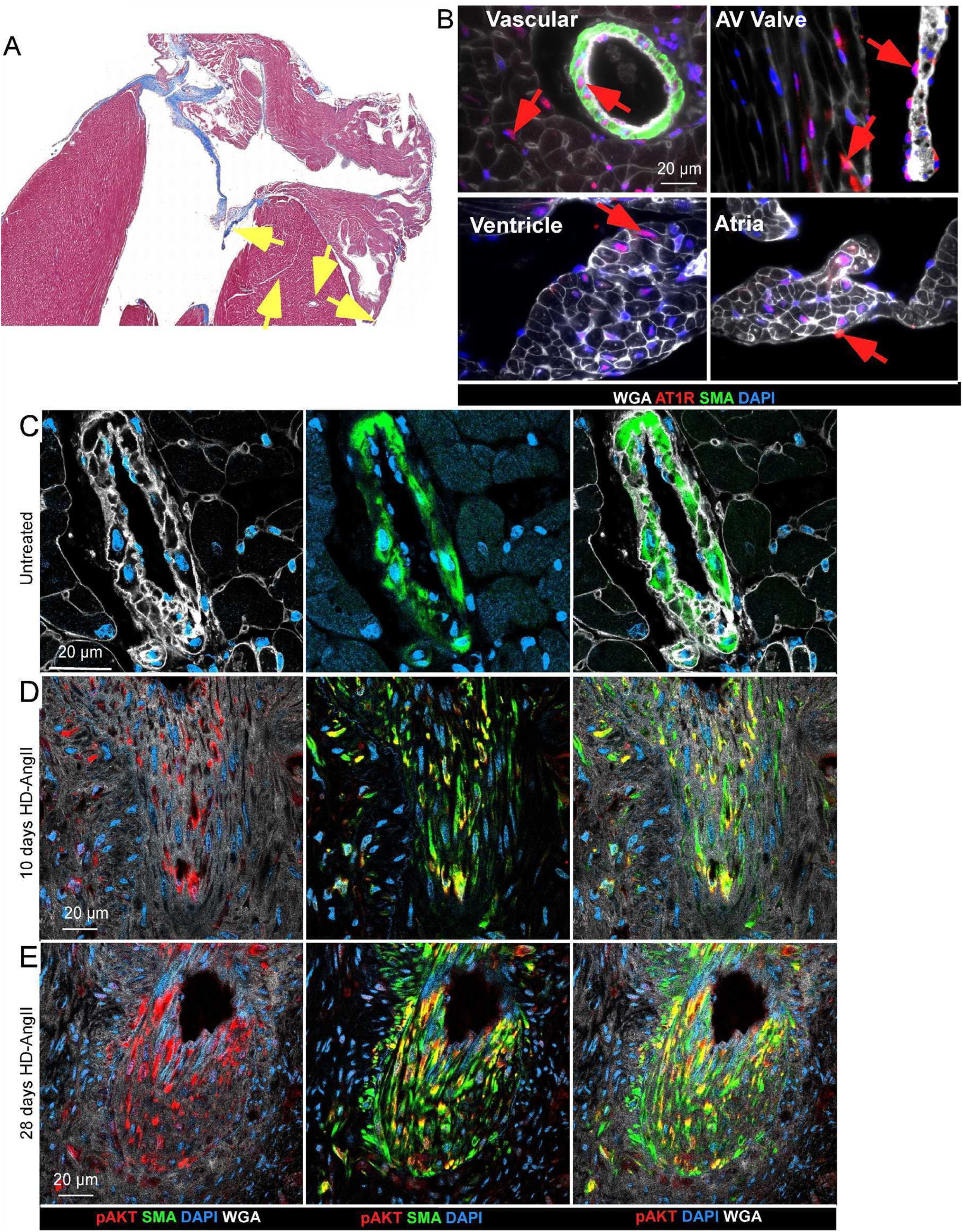
AT1R distribution and pAKT activation in AngII-induced vasculopathy. (A) Representative Masson’s trichrome–stained section of an untreated heart illustrating the region from which the AT1R immunofluorescence insets were obtained (yellow arrows). (B) High magnification views of regions indicated by yellow arrows in A. AT1R expression (red) in smooth muscle cells (SMA; green), endothelial cells, cardiomyocytes (ventricular and atrial), and valvular structures (WGA; white). Yellow arrows highlight regions of interest (vascular, atrioventricular valve, ventricular and atrial insets), and red arrows indicate areas with high AT1R expression. (C-E) Phosphorylated AKT (pAKT; red) in VSMCs (SMA; green) is absent in untreated hearts (C), appears by day 10 of HD-AngII infusion (D), and persists through day 28 (E). pAKT localizes predominantly to cytosolic regions adjacent to the plasma membrane rather than to nuclei (DAPI; blue).

### 2.6 AngII-induced vascular remodeling across systemic vascular beds

We evaluated vascular remodeling across multiple beds, including mesenteric, aortic, skeletal muscle, and renal arteries. Although a comprehensive quantitative analysis was not performed, Masson’s trichrome staining revealed a similar sequence of changes in these vessels, with an initial inflammatory response followed by fibrosis (**Supplemental figure 3**). The aorta appeared relatively resistant to 28 days of HD-AngII infusion, which may reflect differences in the embryologic origin and phenotypic properties of its vascular smooth muscle cells compared with those in coronary and other systemic arteries.

## Discussion

This study demonstrates that hemodynamic and neurohumoral stressors are associated with distinct patterns of myocardial versus vascular remodeling. Pressure-intensive overload stimulus (TAC) is characterized by homogeneous cardiomyocyte hypertrophy with interstitial myocardial fibrosis, whereas AngII-biased neurohumoral stress is associated with a VSMC-centric coronary vasculopathy with perivascular fibrosis and elastic lamina injury. By combining invasive hemodynamics with multiregional histology and lineage tracing, this study distinguishes pressure-intensive from AngII-driven remodeling patterns and highlights vascular pathways that are not readily explained by afterload alone.

#### Discussion Highlights

- <u>Hemodynamic vs. Neurohumoral Stress Effect</u> - distinct remodeling patterns in myocardial interstitium and coronary vasculature
- Differences and similarities in temporal dynamics of myocardial and coronary remodeling
- · <u>Neointimal Formation and VSMC-Driven</u> <u>Vasculopathy</u> - VSMC orchestrate neointima development through centripetal hyperplasia and hypertrophy, progressively narrowing the vascular lumen while inducing mechanical stretch and rupture of EEL/IEL laminae. Endothelial lineage tracing shows that this process occurs beneath an intact endothelial monolayer, with minimal direct endothelial contribution to the neointima.
- <u>Targeted Therapeutic Opportunities</u>: 1) Dual-pathway strategies targeting coronary vasculopathy vs. myocardial fibrosis; 2) Cell-type specific interventions for VSMCs, fibroblasts, endothelial cells, and inflammatory infiltrates
- · <u>Updated Model of VSMC Phenotypic Switching</u> AngII reprograms vascular VSMCs through AT1R → AKT → signaling, driving contractile-to-mesenchymal transition with proliferative, and inflammatory phenotypes.

### Hemodynamic versus neurohumoral stress and compartment-specific remodeling

Historically, hypertension and elevated arterial pressure have been viewed as the primary drivers of vasculopathy, with chronic increases in wall stress leading to endothelial dysfunction, medial hypertrophy, and progressive arterial stiffening. In this framework, vascular injury is largely attributed to long-standing hemodynamic overload and accelerated “vascular aging” [37–39]. Our data refine this paradigm by directly contrasting pressure overload with neurohumoral activation in complementary murine models while acknowledging that these stimuli are biologically intertwined. In the TAC model, marked LV pressure overload produces predominantly myocardial remodeling with cardiomyocyte hypertrophy and interstitial fibrosis and relatively modest large-vessel changes, consistent with prior descriptions of pressure-induced hypertensive remodeling [7–11]. In contrast, AngII/PE and HD-AngII generated lower LV systolic pressures yet produced both myocardial and vascular-dominant remodeling, with extensive perivascular fibrosis and coronary VSMC hyperplasia affecting both the right and left coronary arteries (**Fig. 4A3, 5A2, A2’**). These observations support the concept that TAC produces pressure-dominant remodeling with secondary RAAS activation, whereas AngII/PE and HD-AngII represent AngII-biased states in which neurohumoral signaling preferentially accentuates coronary vascular remodeling rather than simply recapitulating pressure-overload induced myocardial remodeling. Similar pressure-modest, AngII-mediated vascular injury has been reported by others [40, 41] and likely reflects direct AngII effects via AT1R on VSMCs, endothelial, and inflammatory cells [14–16]. Together, these findings support a model in which hemodynamic overload is a major driver of diffuse myocardial fibrosis, while chronic AngII exposure preferentially engages the coronary vasculature and perivascular compartment.

### Temporal dynamics of myocardial versus vascular remodeling

Inflammation emerged early (days 3–10) in both interstitial and perivascular regions and evolved to mature fibrosis by day 28, but the trajectories diverged between compartments. In the myocardium, TAC-induced pressure overload was associated with progressive interstitial ECM deposition and homogeneous hypertrophy, without evidence of elastic fiber involvement (**Supplemental Fig. 2**). In contrast, HD-AngII tightly coupled myocardial remodeling with a VSMC-centric vasculopathy, in which synchronized smooth muscle proliferation, perivascular fibrosis, and neointima formation progressed in parallel with regional, heterogeneous cardiomyocyte hypertrophy and interstitial matrix expansion. These temporal and topographic differences highlight compartment-specific therapeutic windows, suggesting that early anti-inflammatory or anti-proliferative strategies may be required to prevent AngII-driven vasculopathy, whereas SGLT2 inhibitors and aldosterone antagonists primarily improve cardiomyocyte relaxation and reduce myocardial fibrosis, with limited direct vascular effects.

### Vascular remodeling: endothelium, neointima, elastic lamina, and adventitia

In many vascular diseases, endothelial injury is considered the initiating event, leading to barrier disruption [23], increased permeability, immune cell recruitment, and impaired nitric oxide–mediated vasodilation [22, 42]. Current therapeutic strategies, including anticoagulation [43], anti-inflammatory agents [44], and lipid-lowering therapies [45], are largely built around this endothelium-first paradigm. Notably, our HD-AngII model demonstrates robust coronary remodeling without detectable endothelial damage, endothelial-to-mesenchymal transition (EndMT), or endothelial proliferation contributing to neointima formation consistent with prior observations from our group [27], indicating that AngII-driven vasculopathy can arise primarily from VSMC-centric processes rather than classic endothelial denudation. At the same time, the absence of direct endothelial functional readouts in this study means that we cannot exclude important contributions of endothelial dysfunction to the overall phenotype. However, endothelial lineage-tracing data indicate that AngII-driven coronary vasculopathy in this model develops in the setting of a structurally preserved endothelial monolayer and without appreciable direct endothelial contribution to the neointima (Fig. 8). The absence of CDH5-CreER–lineage traced cells within the neointimal compartment argues against a major role for EndMT or endothelial migration as a source of lesion-forming cells, at least under the conditions studied. Together with the intact luminal endothelium, these findings support a model in which coronary neointima formation in AngII-treated mice is driven predominantly by smooth muscle (or SMC-like) lineage cells undergoing phenotypic remodeling, with ECs acting more as stable paracrine regulators and barrier elements rather than as direct structural contributors to the neointima.

The neointima in this setting reflects a VSMC-dominated lesion rather than a mixed, endothelium-initiated response [46, 47]. Lineage tracing (**Fig. 8**) confirms VSMCs as the predominant contributors to neointima formation in our HD-AngII model. We identify multiple VSMC phenotypes—hypertrophic, mesenchymal/undifferentiated, and macrophage-like—that build a non-compacted medial layer and centripetal neointimal expansion. This inward growth narrows the vascular lumen, impairs blood flow, and mechanically stresses the IEL/EEL, creating an unstable vascular wall prone to rupture. Current vasodilator therapies (e.g., calcium channel blockers, cGMP-elevating agents, prostacyclin, α-blockers, and calcium sensitizers) mainly target contractile tone in hypertrophic VSMCs but do not address proliferative, mesenchymal, or inflammatory phenotypes. Agents that modulate VSMC plasticity, such as mTOR inhibitors, partially address this gap by restraining mesenchymal transition and proliferation, yet the spectrum of VSMC states observed in our model underscores the need for broader phenotype-targeted approaches. Our data suggest that stage-specific inhibition of early proliferative and phenotypically switched VSMCs, rather than late-stage fibrosis alone, may offer the most effective window to stabilize vascular architecture and prevent ischemia.

The internal and external elastic laminae (IEL/EEL) normally provide structural integrity and elasticity to withstand pulsatile flow, and their disruption is classically associated with aneurysm and advanced atherosclerosis. In our HD-AngII model, we observe IEL rupture with delamination, porous disruption, and centripetal VSMC penetration, accompanied by neo-elastic fiber synthesis forming circumferential rings in the neointima. These features point to three interconnected mechanisms: (1) inward VSMC proliferation that stretches IEL/EEL outward, paradoxically increasing compliance while destabilizing the wall and promoting aneurysm risk; (2) reactivation of an elastic fiber synthesis program in postnatal VSMCs—a capacity typically lost after development [48]—producing disorganized neointimal fibers of uncertain functional quality; and (3) VSMC architectural disruption, with IEL/EEL loss correlating with medial non-compaction and likely impaired cell–cell and cell–matrix adhesion. Together, these changes frame elastic lamina remodeling as both a marker and a mediator of AngII-driven vasculopathy, and they identify restoration of VSMC–matrix architecture as a potential therapeutic goal.

The adventitia and perivascular region also follow a stereotyped pattern in the HD-AngII model: inflammation appears by day 10 and matures to dense fibrosis by day 28 (**Fig. 5**). Neointimal expansion and medial VSMC changes occur on a similar time course, suggesting coordinatedbut currently incompletely defined signaling between the vessel wall layers and the surrounding perivascular matrix. This simultaneity supports a view of vasculopathy as a multicompartment process, in which intimal, medial, and adventitial changes evolve in parallel rather than sequentially.

### VSMC plasticity and intercellular crosstalk

Vascular cells orchestrate disease progression through direct contact, paracrine signaling, and extracellular vesicles [23, 47], and prior work in atherosclerosis and restenosis has highlighted EC–VSMC–immune cell crosstalk as a central driver of remodeling [49, 50]. Within this context, our HD-AngII model adds two features that may be particularly relevant to AngII-related vasculopathy: (1) VSMC non-compaction in the media, which may facilitate phenotype switching and proliferation and thus represents an architectural as well as molecular therapeutic target; and (2) IEL/EEL rupture and stretch driven by centripetal VSMC forces, creating aneurysm-prone vascular fragility even in the absence of extreme pressures. Building on established VSMC plasticity frameworks [28], we define four phenotypic states—contractile, mesenchymal/proliferative, macrophage-like, and hypertrophic—that coexist and shift over time in response to AngII (**Fig. 11)**. This expanded “wheel” positions VSMC plasticity as a central, multi-state driver of vasculopathy and suggests that successful therapies will need to modulate both state transitions and the mechanical context created by the elastic laminae and perivascular matrix.

**Figure 11.**
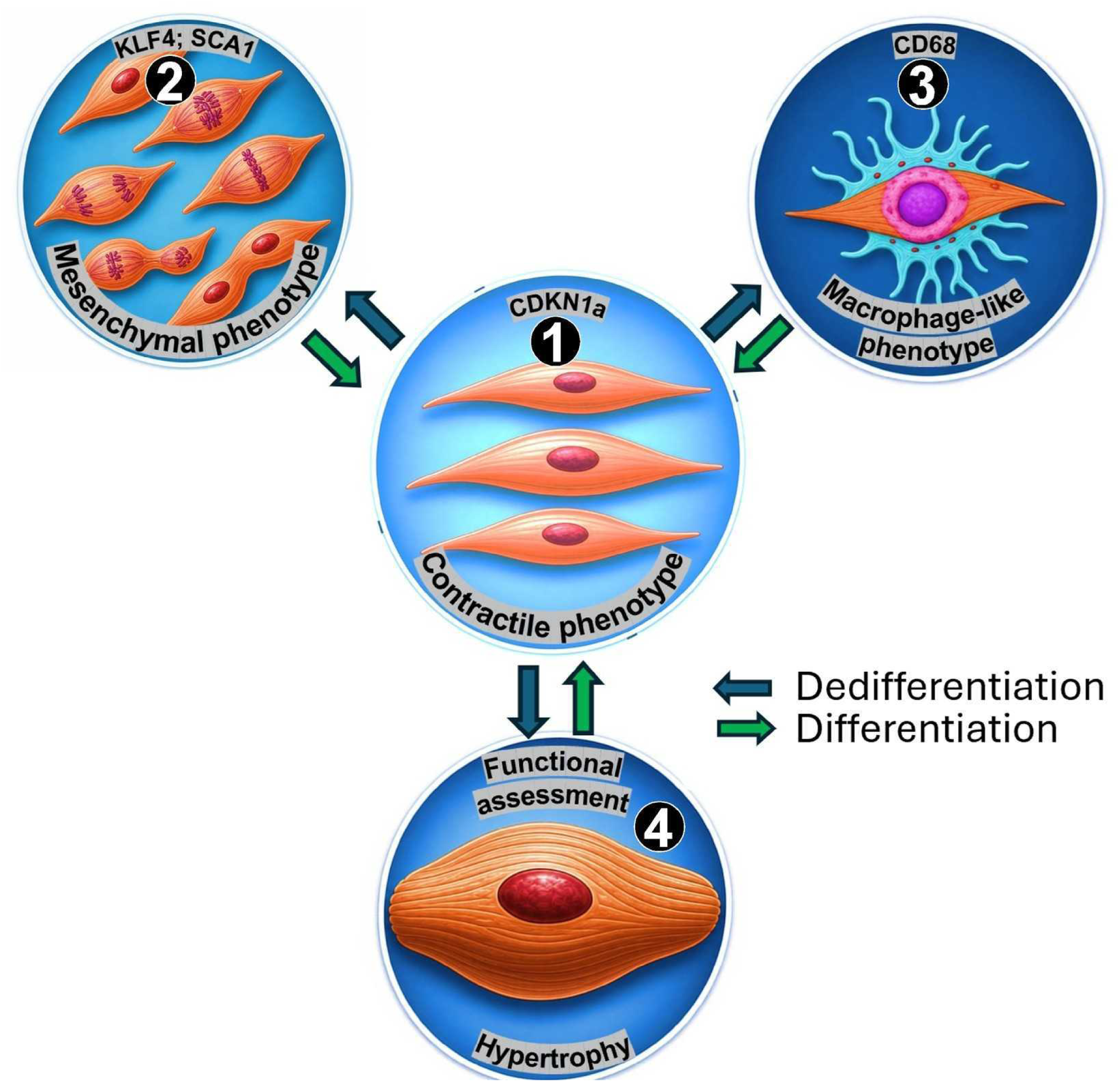
Model of SMC phenotypic switching during AngII infusion. Smooth muscle cells undergo sequential phenotypic transitions: [1] contractile state expressing CDKN1A, [2] undifferentiated state marked by SCA1 and KLF4 expression, [3] macrophage-like state expressing CD68, and [4] hypertrophic phenotype.

### Desmosine as a marker of vascular disease

Desmosine, an elastin-specific cross-linking amino acid, is released during degradation of mature elastic fibers and has been used clinically as a biomarker of elastin breakdown and adverse vascular remodeling. The preferential elevation of serum desmosine in HD-AngII–treated mice with a vascular-predominant phenotype, despite comparable exposure duration and systemic infusion conditions, indicates that AngII-driven vasculopathy is characterized by pronounced elastic lamina injury that is distinct from interstitial myocardial fibrosis (**Fig. 9**). Together with human studies linking circulating desmosine to arterial stiffness [51], coronary disease [52], and adverse outcomes, these findings support the concept that elastin degradation products could serve as fluid biomarkers to identify and track a vascular HFpEF-relevant endotype [31].

### Neurohumoral systemic vasculature stress and vascular remodeling

Beyond the coronary circulation, HD-AngII also induces remodeling in mesenteric, renal, and skeletal muscle arteries, although smooth muscle changes are less pronounced than in coronary vessels (**Supplemental Fig. 3**). This vascular bed heterogeneity likely reflects differences in VSMC embryonic origin and AT1R signaling thresholds, reinforcing the view of the systemic vasculature as a collection of organ-specific phenotypes rather than a uniform compartment [53–55]. Despite these regional differences, HD-AngII produces conserved neointimal pathology across beds, indicating that this model can serve as a platform for studying shared AngII-driven vasculopathies and for testing VSMC-targeted therapies in diverse vascular territories [56].

### Study limitations

**Incomplete stimulus spectrum.** While our models emphasize AngII-biased neurohumoral stress and TAC-induced hemodynamic overload, clinical coronary vasculopathy involves additional factors including metabolic derangements, microvascular dysfunction, and cardiorenal/hepatocardiac syndromes. Nevertheless, we posit that coronary remodeling exhibits conserved temporal progression and histologic hallmarks such as SMC proliferation, medial non-compaction with EEL/IEL instability, and perivascular fibrosis across diverse etiologies. **Stimulus confounding.** Complete isolation of individual stimuli proves challenging. AngII exerts direct neurohumoral actions and indirect hemodynamic effects via vasoconstriction and hypertension, whereas TAC primarily provokes pressure overload with secondary RAAS activation driven by renal hypoperfusion. This secondary RAAS activation could, in principle, be mitigated with ACE inhibitors or ARBs, although such interventions may increase mortality in severe pressure-overload models. These interactions underscore the interdependence of pathologic mechanisms in vivo and motivate our use of “pressure-intensive” and “AngII-biased” rather than purely hemodynamic versus purely neurohumoral terminology. **Phenotypic classification limitations.** Myocardial sampling may miss focal remodeling regions, introducing classification subjectivity between vascular and interstitial phenotypes. We mitigated this through standardized mid-papillary short-axis sections plus four-chamber axial views, though incomplete coverage remains an inherent constraint of tissue-based phenotyping. **Finally,** we did not directly assess functional coronary endpoints (such as coronary flow, flow reserve, or microvascular resistance), which limits our ability to fully quantify the hemodynamic consequences of the observed vasculopathy.

## Conclusion

This study defines distinct yet overlapping drivers of myocardial versus coronary remodeling: hemodynamic stress (TAC) predominantly induces myocardial remodeling characterized by interstitial fibrosis and relatively homogeneous cardiomyocyte hypertrophy, whereas AngII-driven neurohumoral activation is associated with combined myocardial and vascular remodeling with a clear bias toward perivascular fibrosis, elastic lamina injury, and VSMC-centric vasculopathy. These complementary paradigms, with their vascular compartment-specific pathways and temporal dynamics, define AngII-associated vascular remodeling programs that are pertinent to vasculopathies seen in vascular HFpEF and in conditions such as chronic kidney disease and renovascular hypertension. They also provide a conceptual basis for precision therapies targeting AngII-associated vasculopathies, suggesting that optimal treatment will pair myocardial unloading with targeted modulation of VSMC plasticity, elastic lamina integrity, and perivascular fibrosis.

## Methods

### Mice

Mice were housed in a pathogen-free facility and handled in accordance with standard use protocols, animal welfare regulations, and the *NIH Guide for the Care and Use of Laboratory Animals*. Mice were housed with a 12-hour day-night cycle in a temperature (22±1°C) and humidity (55±5%) controlled room. Mice were allowed free access to water and a standard laboratory mouse diet (PicoLab^®^ Rodent Diet 20, #007688). All protocols were approved by the Washington University Animal Studies Committee. All mice were maintained on a C57BL/6J; 129X1 mixed or F1 hybrid genetic background. Both male and female mice were used in these studies with approximately equal distribution.

### Pharmacological induction of vasculopathy

Eight- to ten-week-old mice were implanted with an osmotic mini-pump (Alzet, 200 μl, #2004) to allow subcutaneous infusion of angiotensin II and phenylephrine (AngII/PE) for 28 days as described [27]. Osmotic minipumps were programmed to deliver angiotensin II (1.5 μg/g/day) and phenylephrine HCl (50 μg/g/day) [27, 57–59] or high dose AngII (3 µg/g/day). For controls, cohorts of mice were implanted with saline filled mini pumps. Loaded minipumps were primed in 0.9% NaCl for 24 hr at 37⁰C prior to implantation. Mice were randomly selected and assigned to different groups: control mice (either no pump implantation or implantation of a pump filled with saline), and mice implanted with an infusion pump with AngII/PE or high HD-AngII.

### Transverse aortic constriction (TAC)

Eight- to ten-week-old mice underwent minimally invasive TAC with a severe degree of constriction (27G spacer; 0.41 mm OD) for 4 weeks, alongside sham-operated controls [60]. TAC was performed with a target of 8 mice per group, based on an anticipated minimum survival of 80% from prior studies. All instruments were sterilized in a glass bead sterilizer before surgery, and aseptic technique was maintained throughout.

Mice were weighed, then anesthetized with 2.5% isoflurane in 100% oxygen via nose cone; body temperature was maintained at 37 ± 0.5 °C using a heating pad with rectal probe monitoring. Anesthesia was maintained at 2.0–2.5% isoflurane, and surgical depth was confirmed by absence of the toe-pinch reflex. Buprenorphine SR-LAB (1 mg/kg, ZooPharm, USA) was administered subcutaneously for analgesia. The left anterior and lateral thorax was shaved, prepped with three alternating povidone-iodine and alcohol scrubs, and draped sterilely.

Mice were intubated with a 24G angiocatheter and mechanically ventilated (Model 687, Harvard Apparatus) at 80–90 breaths/min, tidal volume 0.2–0.3 mL. A left thoracotomy (∼1 cm) was performed; pectoralis and intercostal muscles were bluntly dissected, and the ribs retracted. After retracting the thymus, the transverse aorta (between the brachiocephalic and left common carotid arteries) was visualized and carefully isolated. A 7-0 nylon ligature was passed around the aorta and loosely tied; a pre-sterilized blunt 27G needle was placed within the loop, the knot tightened to constrict the aorta around the needle, and the needle removed.

The chest wall (ribs, intercostal, and pectoralis muscles) was closed with running 6-0 absorbable sutures, and the skin with 6-0 nylon. Mice were weaned from the ventilator, extubated, and anesthesia discontinued. Once sternally recumbent, animals recovered under a warming lamp and were monitored closely during the perioperative period.

### Hemodynamic Analysis

Under general anesthesia (isoflurane) and full ventilatory support, retrograde catheterization was performed with a 1-Fr high-fidelity micromanometer pressure catheter (SciSense Advantage System, London, ON, Canada). Systolic and diastolic blood pressure (BP) were recorded at 10, and 28 days in the right carotid artery and analyzed with SciSense software. After advancing the catheter into the LV, LV systolic and diastolic pressures were recorded at 28 days and analyzed with SciSense software.

### Histology and morphometry

Cardiomyocyte cross-sectional area, capillary density, and small-vessel smooth muscle actin were quantified on 6-µm paraffin sections stained with wheat germ agglutinin (WGA), as previously described [27, 61–63]. Interstitial and perivascular cardiac fibrosis were assessed on Masson’s trichrome–stained sections. Regions of interest (ROIs) comprised whole left ventricular (LV) coronal sections at the level of the papillary muscles and included all four cardiac chambers (RA, LA, RV, LV).

Cardiac remodeling patterns (interstitial, perivascular, mixed, and vascular) were classified on Masson’s trichrome–stained coronal sections encompassing the majority of the heart by observers blinded to treatment group, using predefined criteria based on the predominant compartmental localization of inflammation/fibrosis.

Images were analyzed in ImageJ using color thresholding to segment blue collagen (fibrosis) from red myocardium. Interstitial fibrosis was calculated as the fibrotic area excluding perivascular regions, expressed as a percentage of the total myocardial area in the coronal section. Perivascular fibrosis was quantified as the area of collagen surrounding intramyocardial vessels, expressed (i) as a percentage of the total myocardial area in the section and (ii) normalized to the total vessel area, including neointima and tunica media [64, 65].

Vasculopathy was evaluated by three complementary metrics. (1) <u>VSMC</u> <u>compactness</u> (combined neointimal and medial VSMCs) was defined as the summed 2D area of VSMC profiles divided by the combined area of the neointima (between endothelium and internal elastic lamina) and tunica media (between internal and external elastic laminae), providing a measure of cellular packing. (2) <u>Neo-elastic lamellae</u> were quantified as the number of newly formed elastic rings within the intima–media region, excluding the native internal and external elastic laminae. (3) <u>Wall thickening</u> was measured as neointimal and medial areas expressed relative to the total vessel area, including the vascular lumen, consistent with standard neointimal morphometry [66, 67].

Desmosine and AngII concentrations were quantified using commercially available ELISA kits (Cusabio; desmosine: CSB-E14196m, AngII: CSB-E04495m) according to the manufacturer’s instructions.

### Immunofluorescence

Histological sections (6 μm) were prepared from paraffin-embedded tissues. Sections were deparaffinized and rehydrated. Antigen retrieval was performed using a pressure cooker and citrate buffer (pH 6.0). Tissues were blocked with 5% goat or donkey serum. For immunofluorescence, the following primary antibodies were used: Collagen 1 (Abcam 1:100, ab 138492, Lot GR3370247-5), αSMA (1:200, Dako North America, M0851), PECAM1 (CD31, 1:50, Dianova, Dia310), CD45 (1:100, Bioscience, Lot 553380), Mac-2 (1:100, Invitrogen, Bioscience, ref 14-5301-85, Lot 2153405), Ly6g (1:100, Abcam, ab25377; Lot GR3443546-1). Antibodies were added to blocking buffer and slides were incubated overnight at 4°C. Slides were washed with PBS and Alexa Fluor-conjugated secondary antibodies (1:200, ThermoFisher) were added for 30 min at room temp. Immunofluorescent imaging was performed using a Zeiss Apotome II and image processing was performed using Zeiss Axioplan software and ImageJ. Digital scanning of whole slides was performed using a Zeiss Axio Scan.Z1.

### Statistical Analysis

Data was analyzed with GraphPad Prism software. For quantification of biological measures, two samples/conditions were analyzed using the Student’s t-test, shown as mean ± SEM; three or more samples/conditions were analyzed using one-way analysis of variance (ANOVA) with post hoc Tukey’s multiple comparisons test. The data are reported as the means ± SEM, and changes with P values less than 0.05 were considered to be statistically significant.

## Legends

**Supplemental Figure 1.**
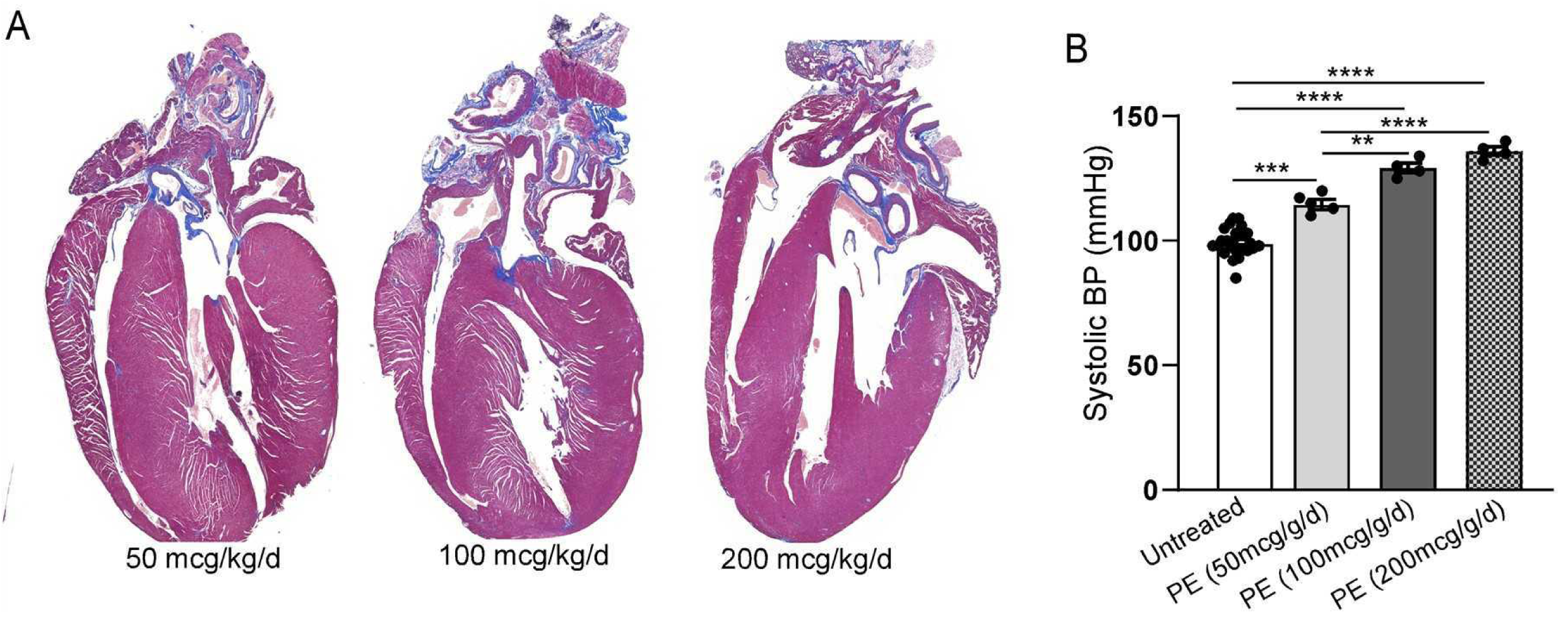
Histologic and hemodynamic effects of graded phenylephrine infusion. Retrograde LV catheterization–derived systolic pressures in mice infused with phenylephrine at (A) 50 μg/g/day, (B) 100 μg/g/day, and (C) 200 μg/g/day. Peak systolic pressure reaches approximately 135 mmHg at 100 μg/g/day and plateaus at about 140 mmHg despite further dose escalation to 200 μg/g/day (N > 10 untreated; N = 5 at 50 μg/g/day; N = 4 at 100 and 200 μg/g/day phenylephrine). Statistical significance was defined as: * P<0.05; ** P<0.01, and *** P<0.001. Data was analyzed using two-way ANOVA with multiple comparisons

**Supplemental Figure 2.**
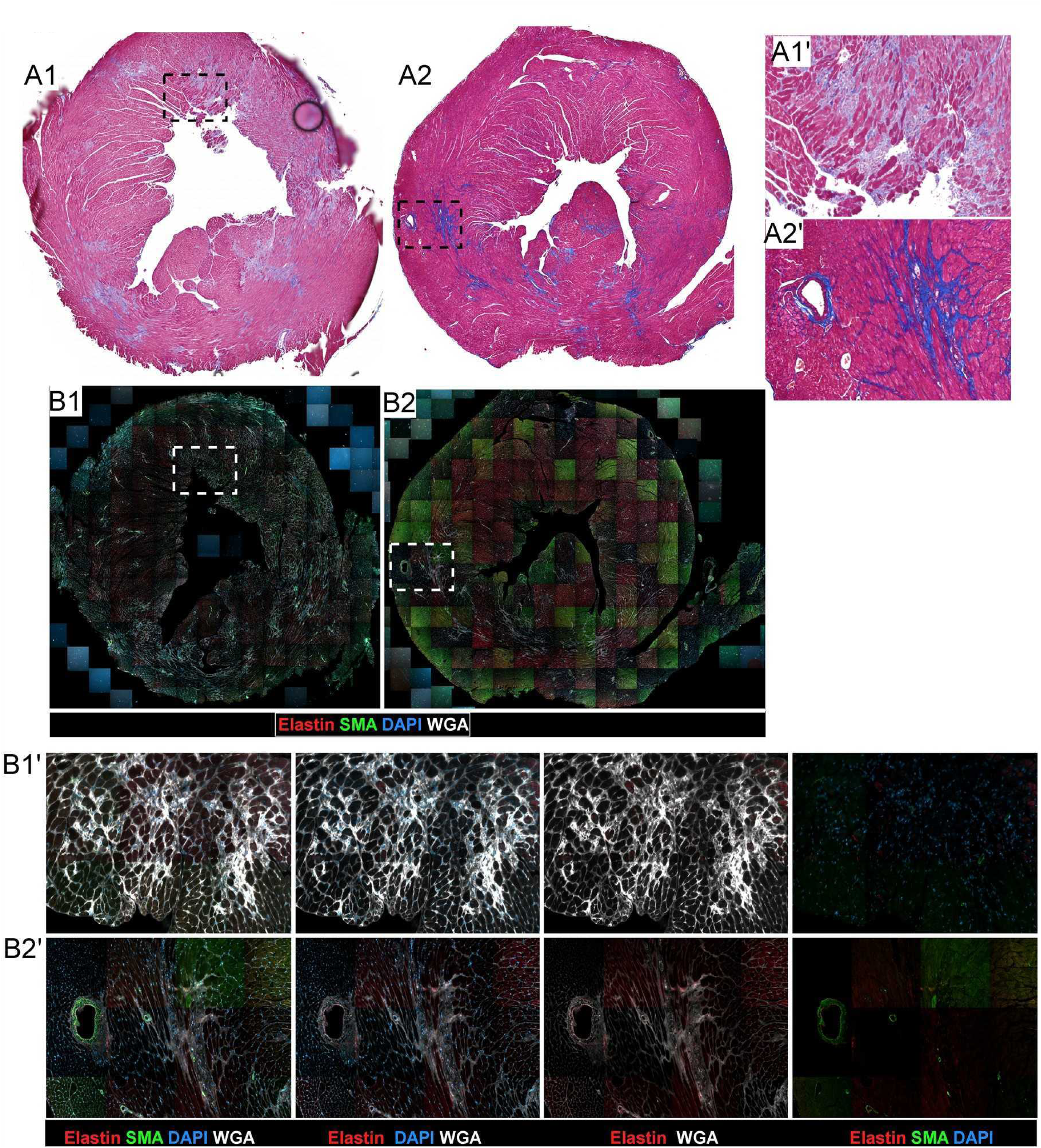
Temporal myocardial ECM remodeling lacking elastic fiber deposition. (A) Masson’s trichrome staining at 7 days (A1) and 28 days (A2) of HD-AngII infusion demonstrates progressive extracellular matrix expansion, with fibrosis becoming more prominent at 28 days (A1′ and A2′ are higher-magnification of boxed regions in (A)). (B) Immunofluorescence at 7 days (B1, high magnification inset B1′) and 28 days (B2, high magnification inset B2′) for WGA (white), elastin (red), SMA (green), and DAPI (blue) shows an absence of elastin accumulation within fibrotic myocardial regions (WGA-positive/blue areas), while elastin signal remains confined to vascular internal and external elastic laminae.

**Supplemental Figure 3.**
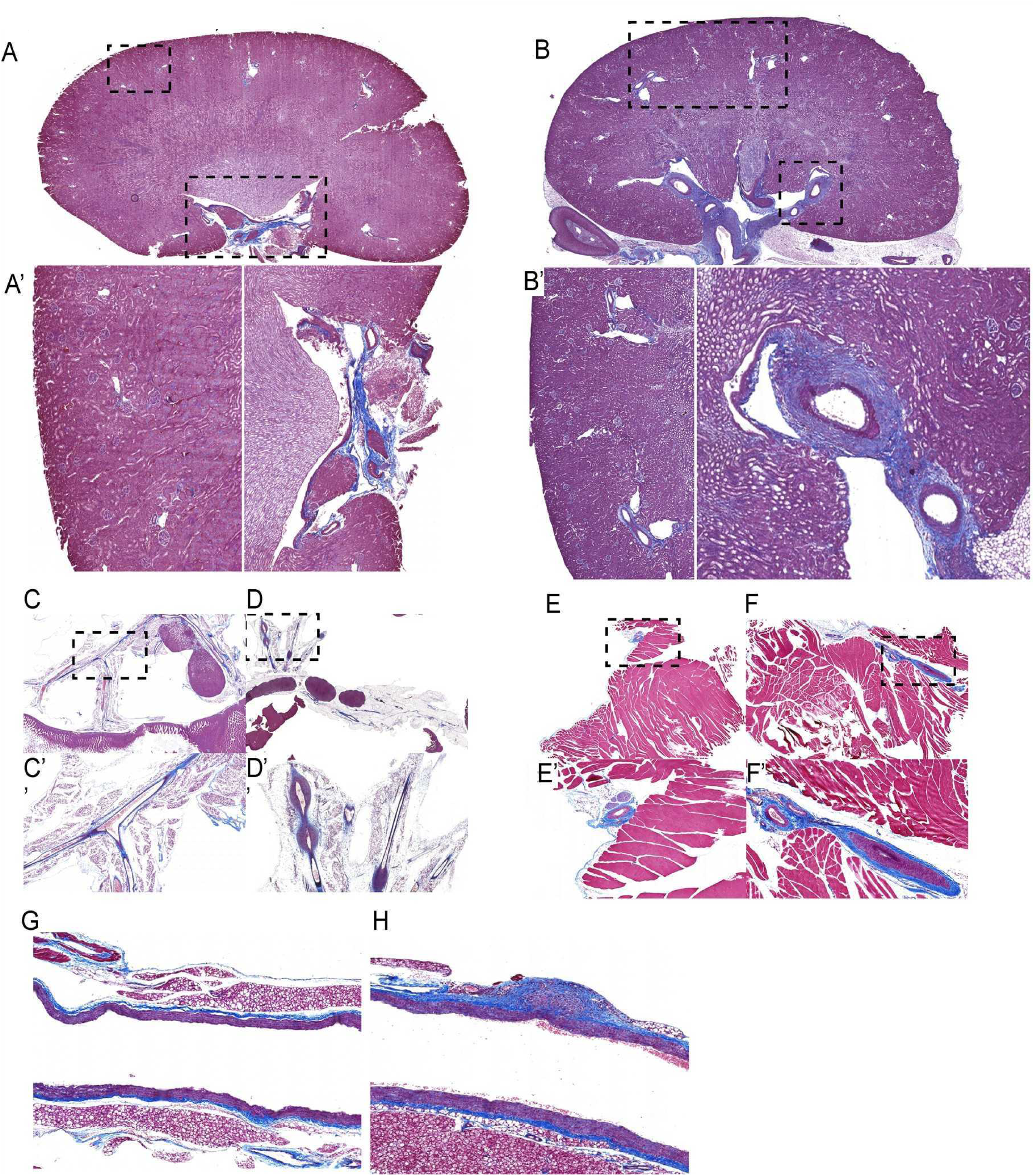
Systemic vascular remodeling across different organ vascular beds. Masson’s trichrome staining of (A) renal vasculature, untreated; A′ and A′′, high-magnification insets of boxed regions in A; (B) renal vasculature after 28 days of HD-AngII infusion; B′ and B′′, high-magnification insets of boxed regions in B; (C) mesenteric second-order branches, untreated; C′, high-magnification inset of boxed region in C; (D) mesenteric branches after 28 days of HD-AngII infusion; D′, high-magnification inset of boxed region in D; (E) skeletal muscle, untreated; E′, high-magnification inset of boxed region in E; (F) skeletal muscle after 28 days of HD-AngII infusion; F′, high-magnification inset of boxed region in F; (G) aorta, untreated; and (H) aorta after 28 days of HD-AngII infusion.

**Supplemental Videos 1 & 2. Dynamic imaging of coronary artery remodeling in the HD-AngII model.**

Video 1. 2D z-stack showing longitudinal views of untreated (1A) and HD-AngII-treated for 10 days (1B) coronary arteries. Video shows a compact, single SMC layer (red) in controls versus IEL rupture with protruding/delaminating SMCs in HD-AngII-treated vessels.

Video 2. 3D reconstructions of IEL/EEL demonstrate intact porous laminae in untreated arteries (2A) and pronounced IEL rupture in 10 days HD-AngII-treated samples (2B).

